# When to learn from elders or peers: accessibility-knowledgeability trade-offs explain variation in age-biased social learning

**DOI:** 10.64898/2026.06.16.732531

**Authors:** Ludovic Maisonneuve, Laurent Lehmann

**Affiliations:** Department of Ecology and Evolution, University of Lausanne, 1015 Lausanne, Switzerland; Department of Social and Behavioral Sciences, Toulouse School of Economics, Université Toulouse Capitole, Toulouse, France

**Keywords:** social learning, life-history, cultural transmission, evolutionary theory

## Abstract

In many animal species, individuals acquire knowledge from others that enhances their survival and reproduction. However, among the many available cultural exemplars, not all provide reliable information. Consequently, individuals tend to choose their exemplars selectively. One widespread pattern is a preference for older individuals, who may have accumulated valuable knowledge through life. Yet empirical studies also show that individuals frequently learn from age peers, suggesting that copying elders is not universally optimal. The ecological and social conditions that favor learning from elders rather than peers, therefore, remain unclear. Here, we investigate the evolutionary drivers of age-biased exemplar choice in age-structured populations where individuals accumulate knowledge over their lifespan. We develop a continuous-age model that captures the coevolution of exemplar age choice and age-specific investments in social learning, individual learning, and the use of acquired knowledge for energy extraction. We show that selection promotes a progressive shift from social to individual learning and from learning to energy extraction with age. Exemplar age choice evolves through a trade-off between knowledgeable and accessible individuals. Young learners therefore favor relatively young exemplars because they are common and still provide substantial novel knowledge, although older individuals remain more knowledgeable. As individuals age, encountering exemplars with substantially novel knowledge becomes increasingly difficult. Consequently, as they age, individuals are expected to shift toward learning from older individuals, who possess more knowledge. Population, environment, and knowledge characteristics can shift this balance, generating a wide range of strategies from learning primarily from peers to consistently targeting the oldest individuals. In particular, learning from age peers is favored by strong within-cohort interactions, high mortality, or high encounter rates. It is also favored in unstable environments with rapid knowledge loss and when knowledge is easily acquired, transmitted, and is bounded.

## 1 Introduction

In many animal species, individuals acquire knowledge from others that enhances their survival and reproduction. However, among the many available cultural exemplars, not all provide the same amount of reliable information (Ralphs et al., 1994; Rieucau and Giraldeau, 2011; Nöbel et al., 2018; Avarguès-Weber et al., 2018; Jackson et al., 2023). Consequently, individuals tend to choose their exemplars selectively (Kendal et al., 2018), for example, by relying on cues of success (e.g., higher performance, Wilkinson, 1992; Kendal et al., 2015; Coelho et al., 2015; Canteloup et al., 2021; Garcia-Nisa et al., 2023, greater physical condition, Duffy et al., 2009 or higher fecundity, Danchin et al., 1998; Doligez et al., 2002; Sarin and Dukas, 2009; Forsman and Seppänen, 2011; Pasqualone and Davis, 2011; Loukola et al., 2013), or by adopting knowledge exhibited by the majority of the population (Danchin et al., 2018). These exemplar choice rules are themselves behavioral traits subject to natural selection. The evolution of exemplar choice involves a feedback loop: individual exemplar choices shape the distribution of knowledge in a population, which in turn modifies the knowledge socially available and the selective pressures on those choices (Lumsden and Wilson, 1980). For instance, when conformity becomes common, it can maintain knowledge that has turned maladaptive after environmental change, as conformists continue to acquire what remains in the majority, thereby reducing conformity’s selective advantage (Henrich and Boyd, 1998; Nakahashi, 2007; McElreath et al., 2008; Perreault et al., 2012). Mathematical models have proven effective in formalizing these interactions between individual choices and population knowledge, and in highlighting feedback effects.

One widespread form of exemplar choice is a preference for older exemplars. Numerous empirical studies show that individuals preferentially acquire or retain information from older conspecifics, including in humans (Rakoczy et al., 2010; Seehagen and Herbert, 2011; Henrich and Broesch, 2011; Wood et al., 2012; Molleman et al., 2019; Bond and Gaoue, 2020; Jang and Redhead, 2025), non-human primates (Coelho et al., 2015; Barrett et al., 2017; Tan et al., 2018; Shorland et al., 2019; van Leeuwen and Hoppitt, 2023; Kukofka et al., 2025; Nodé-Langlois et al., 2025), mice (Choleris et al., 1997), fishes (Dugatkin and Godin, 1993; Amlacher and Dugatkin, 2005), and in meta-analyses across animal species (Camacho-Alpízar and Guillette, 2023). This preference can be advantageous because older individuals may have accumulated greater knowledge over time. For instance, chimpanzees have been shown to master increasingly complex termite-gathering skills as they age (Musgrave et al., 2021), humans in a small-scale society on Pemba Island, Tanzania, have been shown to accumulate substantial ecological knowledge across their lifespan (Pretelli et al., 2022), and Ache hunters improve their ability to locate and capture prey over several decades (Walker et al., 2002).

Other studies, however, reveal that individuals often exchange social information with peers of the same age (e.g., in humans, Ryalls et al., 2000; VanderBorght and Jaswal, 2009; Shutts et al., 2010; Jang et al., 2024, non-human primates, Kawai, 1965, birds, Cornell et al., 2011; Derégnaucourt and Gahr, 2013; Honarmand et al., 2015), and some research finds no detectable age bias in exemplar choice (e.g., in primates, Canteloup et al., 2020). This pattern suggests that preferentially copying older individuals is not universally optimal, but instead contingent on ecological and social conditions. Despite extensive empirical work, however, the evolution of age-biased exemplar choice remains largely unexplored theoretically. A notable exception is the model of Deffner and McElreath (2022), which shows that both “copy-the-old” and “copy-the-young” strategies can be favored under different conditions. In particular, a bias toward younger exemplars can evolve in unstable environments, where young individuals are more likely to hold up-to-date information (Deffner and McElreath, 2022). However, this prediction follows from a fixed learning schedule in which adults cease learning and cannot update their knowledge after environmental change. Allowing learning schedules to evolve may qualitatively alter this result, because unstable environments can instead select for continued adult learning.

More generally, the adaptive value of learning from older individuals is likely to depend not only on the information they possess but also on their availability as social learning exemplars. Although older individuals may accumulate greater knowledge over time, they are often rarer and encountered less frequently than younger individuals. This mechanism can favor peer learning even when older individuals remain more knowledgeable, without requiring environmental change to make younger individuals better informed. Understanding how selection balances this trade-off between exemplar knowledgeability and availability may therefore be key to explaining variation in age-biased exemplar choice.

Here, we study the evolution of exemplar choice and ask how selection can favor strategies ranging from peer learning to preferential learning from the oldest individuals. To this end, we develop a continuous-age model in which exemplar age choice coevolves with age-specific investments in social learning, individual learning, and energy extraction. Rather than dividing the lifespan into fixed learning stages, the model allows individuals to adjust their allocation among these activities at every age. We represent knowledge as a quantitative state that accumulates over life, thereby accounting for gradual variation in the value of exemplars of different ages.

The model explicitly accounts for exemplar accessibility through two features. First, we introduce structure in social interactions, such that individuals disproportionately encounter others of similar age (a pattern documented across a wide range of taxa, including humans, McPherson et al., 2001, non-human primates, Carter et al., 2015, marmots, Wey and Blumstein, 2010, sea lions, Wolf et al., 2007, and birds, Franks et al., 2020; Aplin et al., 2021). Second, we incorporate explicit demographic structure, allowing population age composition to shape the pool of available cultural exemplars. In many species, juveniles outnumber adults (e.g., Hill et al., 2001; Fang et al., 2022), which may increase the likelihood of encountering younger individuals independently of any intrinsic copying preference. Together, these two forms of accessibility can generate an advantage for peer learning: peers may be encountered more frequently than older individuals and, despite being less knowledgeable on average, may still possess substantial knowledge that is novel to the learner. Using evolutionary invasion analysis coupled with optimal control theory approaches, we investigate how population, environment, and knowledge characteristics influence the co-evolution of age-biased exemplar choice and learning schedules.

## 2 Methods

### 2.1 Biological assumptions

We consider a large age-structured population in continuous time where individuals reproduce asexually. Each individual in the population accumulates knowledge, treated as a quantitative variable that ultimately increases its capacity to reproduce. We assume that individuals first pass through a juvenile phase, which we do not model explicitly, during which they grow and start accumulating knowledge (see fig. 1 for the model’s overview). Upon reaching maturity, individuals can reproduce and allocate their time among three competing activities: social learning, individual learning, and energy extraction. Three evolving trait schedules (or age-dependent function-valued traits), *ν, s*, and *φ* (Table 1 for a list of symbols), determine how individuals allocate their time across these activities and select their social learning exemplar at each age after maturity. The values *ν*(*a*) and *s*(*a*) of the first two functions specify, respectively, for a learner of age *a*, the fraction of time instantaneously allocated to learning versus energy extraction, and, among the time allocated to learning, the fraction allocated to social versus individual learning. The third function, *φ*, determines the choice of exemplar age such that *φ*(*a*) is the age of the exemplar targeted by a social learner of age *a*. We assume that exemplars are chosen exclusively among mature individuals.

**Table 1:** Key symbols and their definitions.

| Table 1: Key symbols and their definitions |  |
| --- | --- |
| Model parameters |  |
| $k_0$ | Knowledge at maturity |
| $\beta$ | Social learning rate |
| $\sigma$ | Selectivity in exemplar age selection |
| $\alpha$ | Sampling rate of beneficial information |
| $K$ | Total environmental knowledge available |
| $\epsilon$ | Knowledge loss rate |
| $\xi$ | Encounter intensity |
| $f$ | Social fluidity |
| $\rho$ | Strength of age-assortative interactions |
| $\mu$ | Mortality rate |
| $c$ | Energy-to-fecundity conversion factor |
| $e_{\min}, e_{\max}$ | Minimum and maximum rates of energy extraction |
| $k_{1/2}$ | Knowledge level at which half of the additional energy extraction from knowledge is gained |
| $\gamma_s, \gamma_i, \gamma_b$ | Returns exponents for social learning, individual learning, and energy extraction |
| Variables for traits |  |
| $\mathbf{u}_\bullet, \mathbf{u}$ | Focal and resident trait schedule vector |
| $\mathbf{u}_\bullet(a) = (\nu_\bullet(a), s_\bullet(a), \varphi_\bullet(a))$ | Fraction of time invested in learning vs. energy extraction, fraction of learning time allocated to social vs. individual learning, and choice of exemplar age at age $a$ |
| $\mathbf{u}^*$ | Numerically identified convergence-stable trait schedule vector |
| Variables for knowledge and its transmission |  |
| $r_{\text{sl}}(a, \varphi_\bullet(a), k_\bullet(a)), r_{\text{il}}(k_\bullet(a))$ | Potential rates of knowledge acquisition under pure social and pure individual learning |
| $A(a, \varphi_\bullet(a))$ | Rate at which a focal accepts a suitable exemplar at age $a$ |
| $P_a(a', \varphi_\bullet(a))$ | Probability that a focal with target exemplar age $\varphi_\bullet(a)$ selects an encountered individual of age $a'$ |
| $k_\bullet(a), k(a)$ | Focal and resident knowledge at age $a$ |
| Variables for reproduction and selection |  |
| $l_\bullet(a), l(a)$ | Probability that a focal or a resident survives to age $a$ |
| $n(a)$ | Equilibrium density of resident of age $a$ |
| $N(\mathbf{u})$ | Equilibrium population size of a resident population with trait schedule $\mathbf{u}$ |
| $D(N)$ | Density-dependent population regulation function |
| $P(\mathbf{u}_\bullet(a), k_\bullet(a))$ | Instantaneous energy extraction rate of a focal with knowledge $k_\bullet(a)$ |
| $b(\mathbf{u}_\bullet(a), k_\bullet(a), N(\mathbf{u}))$ | Instantaneous birth rate at age $a$ of a focal expressing trait $\mathbf{u}_\bullet(a)$, carrying knowledge $k_\bullet(a)$, in a resident population of size $N(\mathbf{u})$ |
| $R(\mathbf{u}_\bullet, \mathbf{u})$ | Basic reproductive number of a focal lineage with trait schedule vector $\mathbf{u}_\bullet$ in a resident population with trait schedule vector $\mathbf{u}$ |
| $S(\mathbf{u}, a)$ | Age-specific selection gradient |

**Figure 1:**
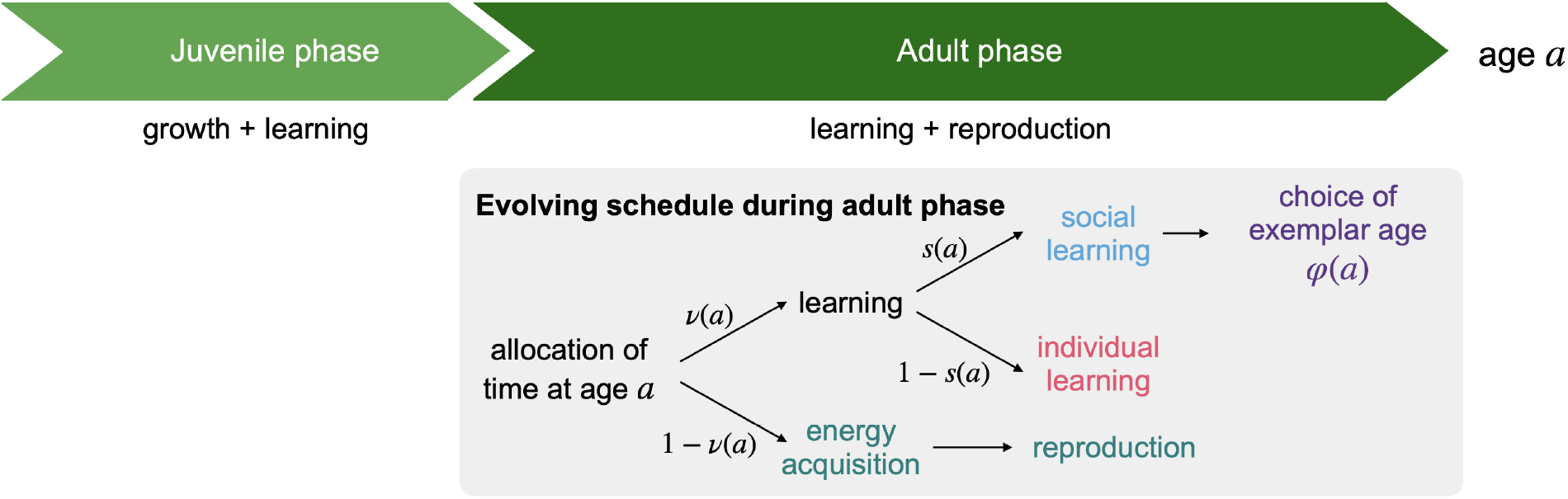
Overview of the model. Individuals first pass through an unmodeled juvenile phase during which they grow and begin accumulating knowledge. After maturity, they reproduce and allocate attention among social learning, individual learning, and energy extraction. Three age-dependent traits, *ν, s*, and *φ*, govern this allocation and the choice of exemplar age. For each age *a, ν*(*a*) specifies the fraction of attention devoted to learning (vs. energy extraction), *s*(*a*) the fraction of learning allocated to social (vs. individual) learning, and *φ*(*a*) the age of the exemplar targeted.

We aim to establish the trait schedules *ν, s*, and *φ* that are favored by natural selection in order to understand how exemplar choice coevolves with learning behavior across the lifetime. To that aim, we carry out an evolutionary invasion stability analysis for quantitative trait schedules assuming that mutations are rare enough to arise in a population monomorphic for some resident schedule (e.g., quantitative trait invasion analysis, Eshel, 1983; Taylor, 1989; Geritz et al., 1998, here applied to cultural evolution, e.g., Aoki et al., 2012; Wakano and Miura, 2014; Mullon and Lehmann, 2017; Maisonneuve et al., 2025; Maisonneuve and Lehmann, 2026). We next detail how the trait schedule ***u***_•_ = (*ν*_•_, *s*_•_, *φ*_•_) of a focal (or representative) individual–and who could bear a mutant or a resident trait–facing a resident population with trait ***u*** = (*ν, s, φ*) influences learning (Section 2.2) and reproduction (Section 2.3).

### 2.2 Learning

Consider a focal individual with trait schedule vector ***u***_•_ = (*ν*_•_, *s*_•_, *φ*_•_). Let *k*_•_(*a*) denote the amount of knowledge held by that individual at age *a* ∈ [0, 1], where ages are scaled so that *a* = 0 corresponds to the age of maturity and *a* = 1 to the maximum possible lifespan (implications of this assumption are discussed in the Discussion). The rate d*k*_•_(*a*)*/*d*a* of change in the focal individual’s knowledge after reaching maturity is assumed to be given by

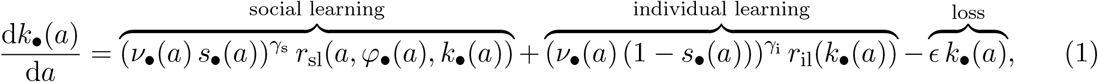

with *k*_•_(0) = *k*_0_, where the parameter *k*_0_ is the amount of knowledge the focal individual has acquired when juvenile, for instance, through learning within the family.

The terms *r*_sl_(*a, φ*_•_(*a*), *k*_•_(*a*)) and *r*_il_(*k*_•_(*a*)) describe the potential rates of knowledge acquisition under pure social learning and pure individual learning, respectively (their explicit forms are given below). Actual rates of knowledge acquisition depend on the time instantaneously allocated to each learning mode through the terms 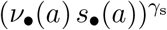 and 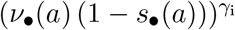, where the exponents *γ*_s_ *<* 1 and *γ*_i_ *<* 1, reflecting diminishing returns to time investment in each learning mode. We further assume that knowledge is lost at a constant rate *ϵ*, capturing both forgetting and obsolescence due to environmental change.

#### 2.2.1 Social learning

We now detail the assumptions underlying the social learning process. At each age *a*, the focal individual seeks a new exemplar of age *φ*_•_(*a*) ∈ [0, 1]. We model this search as a stochastic encounter process. Encounters between the focal at age *a* and an individual of age *a*^*′*^ occur at rate 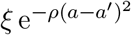, meaning that individuals are more likely to interact with others close to their own age. This could reflect, for example, that individuals of similar ages tend to engage in similar activities. Larger values of *ρ* make interactions more strongly concentrated among similar ages, while *ρ* = 0 corresponds to random interactions across ages.

After an encounter, the focal individual cannot immediately engage in another interaction, as it remains occupied interacting with the encountered individual, regardless of whether it selects them as a social learning exemplar. We assume that interaction durations follow an exponential distribution with mean 1*/f*, where *f* is a social fluidity parameter. When the focal of age *a* encounters a potential exemplar of age *a*^*′*^, it accepts them with probability

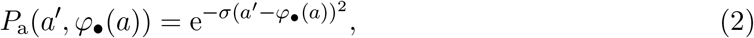

and rejects them with probability 1 − *P*_a_(*a*^*′*^, *φ*_•_(*a*)), where *σ* controls the selectivity of exemplar choice. We assume *σ* to be large, so that the focal accepts only exemplars whose age is close to the target age *φ*_•_(*a*). Higher values of *f* correspond to more fluid social interactions, allowing individuals to encounter others more frequently.

We show in Appendix A.1 that the rate at which the focal of age *a* accepts suitable exemplars can be approximated by

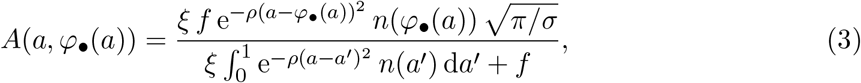

where *n*(*a*) is the equilibrium density of individuals of age *a* in the resident population (see Appendix B for its characterization).

Once the focal accepts an exemplar, the knowledge it can acquire depends on that held by that exemplar. We denote by *k*(*a*) the knowledge held by a resident individual of age *a*, and since the population is large, this is not affected by the focal’s own knowledge.

We view social learning as a foraging process for beneficial pieces of information. Each piece of information held by the exemplar but not by the focal is assimilated independently at rate *β*.

When the time required to handle each piece is negligible, the total rate of acquisition is therefore proportional to the amount of novel knowledge available from the exemplar, *k*(*φ*_•_(*a*)) − *k*_•_(*a*) (Maisonneuve et al., 2025). Thus, the potential rate of knowledge acquisition under pure social learning is

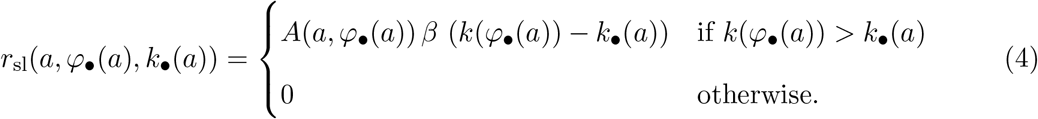

#### 2.2.2 Individual learning

Under the individual learning process, the focal individual is assumed to acquire knowledge at rate

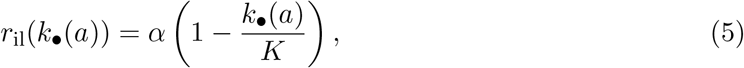

where *α* can be interpreted as the rate at which a naïve individual (i.e., with *k*_•_(*a*) = 0) acquires information, and *K* is the maximum amount of knowledge individuals in the population can potentially acquire (in Appendix A.2 we provide a mechanistic justification for eq. (5) by envisioning individual learning as a process of exploration and retention of beneficial information). The parameter *K* reflects constraints imposed by cognitive capacity and/or the environment. In a population with limited cognitive capacity, individuals can only acquire knowledge pertaining to a few simple environmental challenges relevant to energy extraction (e.g., finding food), yielding a small *K*. In a population with strong cognitive capacity, individuals can acquire and refine knowledge across a wide range of environmental challenges (e.g., processing food, hunting), and the population can potentially accumulate a large amount of knowledge, corresponding to a large *K*. In the limit *K* → ∞, knowledge accumulation becomes potentially unbounded. Importantly, *K* is the maximum amount of knowledge that can potentially be acquired, not the amount reached within one lifespan. The realized knowledge trajectory *k*(*a*) depends on the time allocated to learning, knowledge loss, and intergenerational social transmission.

The set of equations just derived (eqs. 1–5) characterizes knowledge acquisition of a focal individual throughout the lifespan when facing a resident population. When the subscript • is omitted therein in all variables, these equations characterize the knowledge acquisition in the resident population and can be solved for *k*(*a*), so that this quantity gives the equilibrium knowledge of a resident individual of age *a* (though in practice this is a complicated step because knowledge flows asymmetrically across age-classes and depends on the evolving trait schedules, see Appendix B for the full characterization of knowledge dynamics in the resident population). This in turn fully determines the knowledge *k*_•_(*a*) of a focal individual of age *a* (through eqs. 1– 5), and we can now turn to describe how this accumulated knowledge translates into energy extraction and, ultimately, reproductive output.

### 2.3 Reproduction and death

We assume that the instantaneous energy extraction rate of an individual expressing traits ***u***_•_(*a*) and carrying knowledge *k*_•_(*a*) is

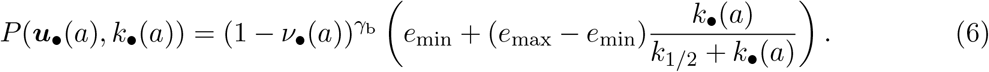

Energy extraction increases with the fraction of time allocated to this activity, 1 − *ν*_•_(*a*) with *γ*_b_ *<* 1 implying diminishing returns. The parameter *e*_min_ is the baseline rate of energy extraction in the absence of knowledge, while *e*_max_ is the maximal energy extraction rate achieved when knowledge is large. The parameter *k*_1*/*2_ is the level of knowledge at which the individual obtains half of the additional energy that can be gained using knowledge (i.e., above the baseline *e*_min_). When knowledge is large, energy extraction approaches *e*_max_.

The difference *e*_max_ − *e*_min_ therefore captures how strongly knowledge can improve energy extraction. Across taxa, ecological knowledge can play an important role in resource acquisition and survival (Swaney et al., 2001; Foley et al., 2008; Thornton and Clutton-Brock, 2011; Brent et al., 2015). Among human foragers, a one-standard-deviation increase in hunting knowledge was associated with roughly 0.8 kg/h more game, on the order of the mean hunting return itself, of 0.98 kg/h (Reyes-García et al., 2016). Experiments in bumblebees show that the slowest-learning colonies collected 40% less nectar than the fastest-learning colonies under natural conditions (Raine and Chittka, 2008). In killer whales, maternal death increased mortality risk in the following year 3.1-fold in sons aged 30 years or less and 8.3-fold in older sons, with the loss of maternal ecological knowledge proposed as one possible mechanism (Foster et al., 2012). These observations suggest that the effect of knowledge on energy extraction can be substantial in some species.

We assume that individuals incur a constant instantaneous maintenance cost *M* at each age. The remaining energy is directly converted into fecundity. Accordingly, for a focal individual expressing traits ***u***_•_(*a*) and carrying knowledge *k*_•_(*a*), the instantaneous birth rate is given by

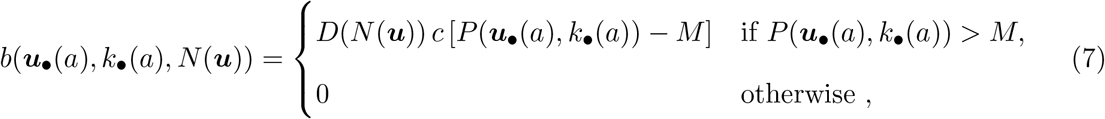

where *c* is a factor converting energy into fecundity and *D*(*N* (***u***)) captures density-dependent regulation, where *N* (***u***) denotes the equilibrium adult population size of the resident population (see Appendix B for its characterization). We assume that a newborn survives the juvenile phase and becomes an adult of age 0 with probability *s*_0_.

We finally assume that individuals face a constant instantaneous mortality rate *µ* when maintenance needs are met, and die instantaneously otherwise.

### 2.4 Evolutionary analyses

Under the above assumptions, the basic reproductive number (e.g., Michod, 1979; Charlesworth, 1980; Inaba, 2017) of an individual with trait ***u***_•_ in an otherwise monomorphic population for trait ***u*** residing at its cultural-demographic equilibrium (see Appendix B) is given by

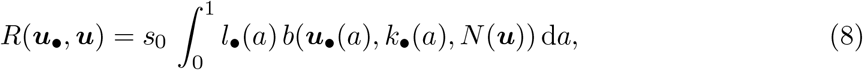

where the term *l*_•_(*a*) = *e*^−*aµ*^ is the probability that an adult with the function-valued trait ***u***_•_ and alive at age 0 survives to age *a*. From standard results on age-dependent branching processes (Mode, 1969, 1971), the lineage of individuals bearing trait ***u***_•_ and descending from a single ancestor with that trait (and necessarily introduced by mutation if this is a new trait value), goes extinct with probability one if *R*(***u***_•_, ***u***) ≥ 1, but has a positive probability of establishing if *R*(***u***_•_, ***u***) *>* 1. Thus *R*(***u***_•_, ***u***) is an appropriate proxy for the invasion fitness of a mutant with trait ***u***_•_ in the resident population ***u*** from which the evolutionary dynamics can be inferred.

In order to understand the direction of selection on trait values expressed at age *a*, we then use the age-specific selection gradient

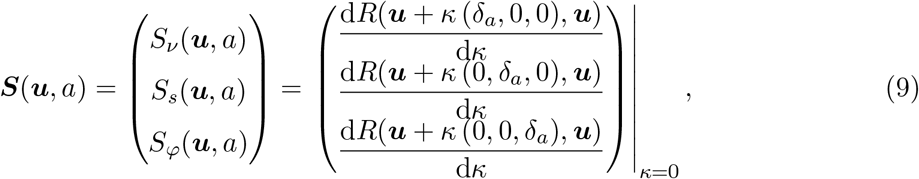

which is a vector of functional derivatives, each component measuring the effect of a small mutational effect in one trait localized at age *a* ∈ [0, 1]. Here, *δ*_*a*_ denotes the Dirac delta function centered at *a* with value *δ*_*a*_(*a*^*′*^) in *a*^*′*^ being zero for all *a*^*′*^≠ *a*. It thus allows to concentrate the (infinitesimal) trait change at *a*, so that the resident schedule ***u*** = (*ν, s, φ*) is unchanged at all other ages (see Appendix C.1 for details).

Because the evolving trait is a function-valued trait, we use optimal control theory (e.g., Bryson and Ho, 1975; Kamien and Schwartz, 2012; Caputo, 2005 for textbook references and e.g., Perrin, 1992; Day and Taylor, 2000; Avila et al., 2021 for applications in evolutionary biology) to get the explicit expressions for the age-specific selection gradient vector, which we then analyze (see Appendix C.2 and Appendix C.4.1, C.4.2 and Section 3.2). Moreover, by iterating the age-specific selection gradient from an initial population without learning, we identify a convergence-stable (e.g., Leimar, 2009) trait schedule vector ***u***\* = (*ν*\*, *s*\*, *φ*\*) (see Appendix C.3). Because ***u***\* is obtained through this iterative procedure, we expect it to be reached by the evolutionary dynamics of the ancestral population. Nevertheless, we cannot guarantee convergence to ***u***\* rather than to another, unidentified convergence-stable trait schedule vector, given the difficulty of fully characterizing the evolutionary dynamics of function-valued traits. Once ***u***\* is identified, we systematically examine whether selection is stabilizing in a population at ***u***\*, favoring a population monomorphic for ***u***\*, or disruptive, promoting polymorphism in trait schedule vectors (see Appendix C.3). In all cases considered in this paper, selection at ***u***\* is found to be stabilizing.

Note that the numerical identification of ***u***\* is computationally intensive, as working with continuous function-valued traits poses a challenging numerical problem that limits the range of parameter values we can feasibly explore. The analytical expressions of the age-specific selection gradients are therefore particularly valuable, as they allow us to assess the robustness of our results across parameter space without relying solely on numerical computation.

## 3 Results

### 3.1 Individuals learn socially first, then individually, before acquiring energy

We first investigate how individuals adjust their learning and energy extraction behavior across the lifespan. To do so, we examine the allocation of time to individual learning, social learning, and energy extraction at the numerically identified convergence-stable trait schedule vector ***u***\*. This analysis reveals that individuals invest primarily in learning early in life and progressively shift investment toward energy extraction with age, allowing knowledge accumulated early to be exploited later to improve energy extraction efficiency (see fig. 2a). Toward the end of life, individuals allocate all their time to energy extraction, which is directly converted into fecundity, as further knowledge accumulation yields no fitness benefits. To assess the robustness of this result, we examine the expression of the age-specific selection gradient for the proportion of time allocated to learning versus energy extraction, *ν*(*a*), which shows that the selective advantage of investing in learning typically declines with age (see Appendix C.4.1), confirming that this life-history pattern is a general feature of the model.

**Figure 2:**
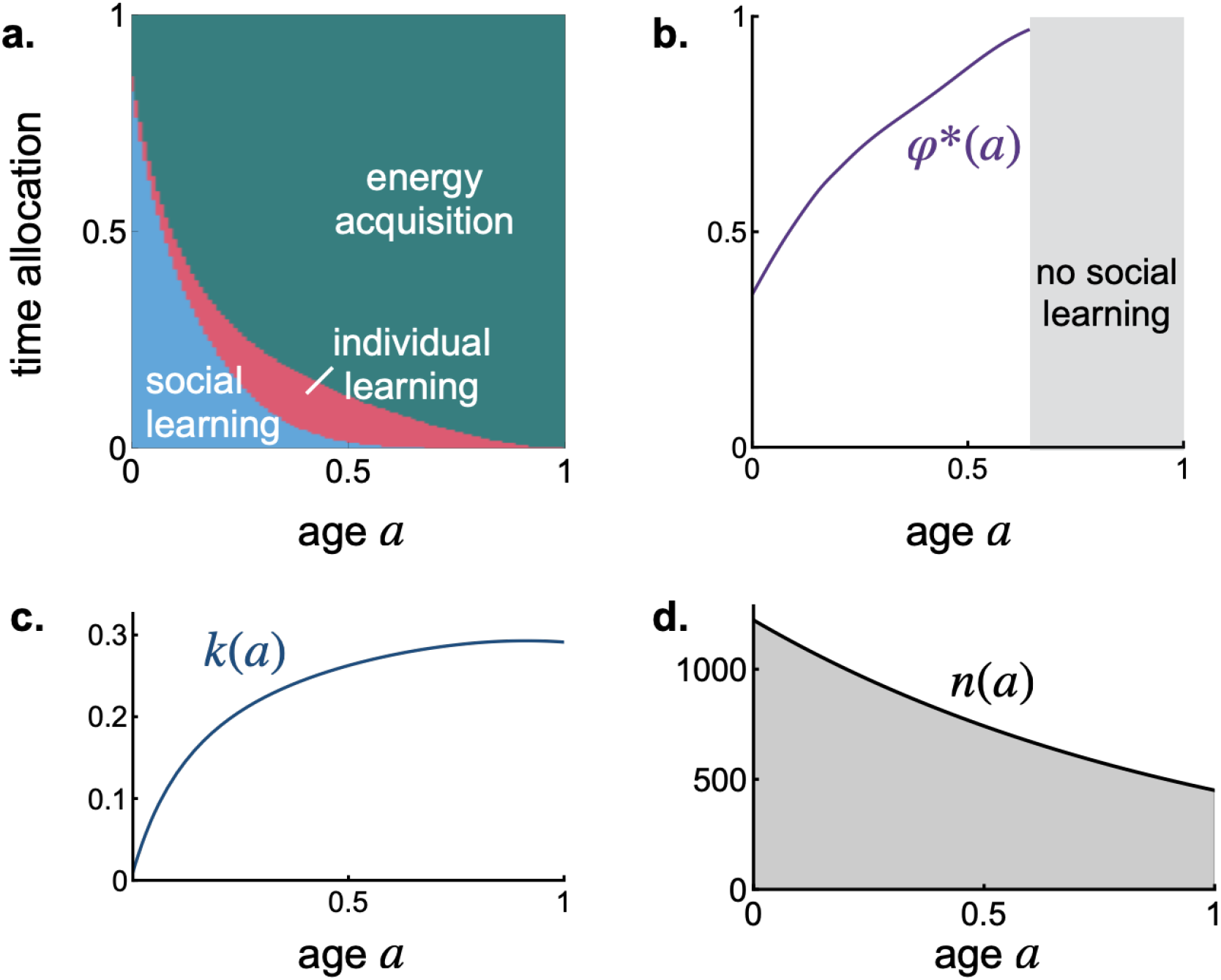
Age-dependent learning behaviors, lifetime knowledge accumulation and age-specific density at the identified convergence-stable trait schedule vector *u*\* = (*ν*\*, *s*\*, *φ*\*). **a** Time allocation schedule at ***u***\*. Colors indicate the proportion of time invested in social learning (blue), individual learning (pink), and energy extraction (teal) at age *a*. At each age *a*, these proportions are given respectively by *ν*\*(*a*)*s*\*(*a*), *ν*\*(*a*)(1 − *s*\*(*a*)), and 1 − *ν*\*(*a*). **b** Choice of exemplar age, *φ*\*(*a*), at age *a* at ***u***\*. **c** Individual knowledge, *k*(*a*), at age *a* at ***u***\*. **d** Age-specific density, *n*(*a*). Parameters are: *ξ* = 1, *f* = 100, *ρ* = 0.01, *σ* = 100, *k*_0_ = 0.01, *α* = 0.5, *β* = 0.6, *ϵ* = 0.1, *K* = 10, *µ* = 1, *e*_min_ = 5, *e*_max_ = 24, *k*_1*/*2_ = 0.2, *M* = 0.01, *γ*_s_ = *γ*_i_ = 0.5, *γ*_b_ = 0.25, *s*_0_ = 0.9, *c* = 10 and *D*(*N*) = 1*/*(1 + 0.1 *N*).

The allocation schedule at ***u***\* further reveals that, among the time allocated to learning, individuals prioritize social learning early in life and gradually shift toward individual learning with age. When individuals are young and have little knowledge, social interactions expose them to many new pieces of information, allowing them to learn quickly from others. As they age and accumulate knowledge, genuinely new information becomes harder to encounter through social interactions, making social learning less effective and encouraging greater investment in individual learning. This result is also robust: analysis of the age-specific selection gradient expression for the proportion of time *s*(*a*) invested in social rather than individual learning, shows that the selective advantage of investing in social learning typically declines with age (see Appendix C.4.2).

### 3.2 Individuals increasingly seek older individuals as they age

We now investigate how selection acts on the age *φ*(*a*) of the exemplar individual each learner seeks at age *a*. Exemplar age choice is shaped by a trade-off between exemplar knowledgeability and accessibility. Targeting older individuals generally provides access to more knowledge, but these individuals are less abundant and, when interactions are strongly age-assortative, less likely to be encountered by younger learners. The age-specific selection pressure formalizes this trade-off:

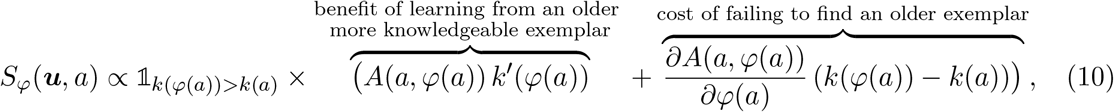

(see Appendix C.4.3 and recall that ∝ means equal up to a constant of proportionality). The first term captures exemplar knowledgeability. The derivative *k*^*′*^(*φ*(*a*)) = d*k*(*φ*(*a*))*/*d*φ*(*a*) captures how targeting older exemplars affects the amount of knowledge available for social learning. Because individuals typically accumulate knowledge over their lifetime, we generally have *k*^*′*^(*φ*(*a*)) *>* 0 for all *φ*(*a*) ∈ [0, 1] (e.g., fig. 2c). Consequently, the term *A*(*a, φ*(*a*)) *k*^*′*^(*φ*(*a*)) is generally positive and favors learning from older, more knowledgeable exemplars.

The second term captures exemplar accessibility. The term *∂A*(*a, φ*(*a*))*/∂φ*(*a*) describes how targeting older exemplars affects the probability of finding a suitable exemplar. In Appendix C.4.4, we show that

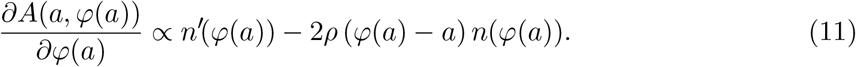

The derivative *n*^*′*^(*φ*(*a*)) describes how the availability of potential exemplars changes with their age. In a population at demographic equilibrium, the density of individuals declines with age, so that *n*^*′*^(*φ*(*a*)) *<* 0 (see Appendix C.4.4 for a proof and fig. 2d). This term is therefore negative and selects targeting younger exemplars. The second term captures the effect of the population’s interaction structure, whereby individuals are more likely to encounter others of similar age, on the probability of finding a suitable exemplar, thereby favoring learning from exemplars of similar age. This second term is generally negative because typically *φ*(*a*) *> a*, as learners can only acquire knowledge from individuals older than themselves who possess greater knowledge. Consequently, this effect also favors targeting younger exemplars.

This decrease in accessibility generates a cost of targeting older individuals. This cost scales with the population-average knowledge difference between targeted exemplars and learners, *k*(*φ*(*a*))−*k*(*a*), because the expected opportunity loss is proportional to the amount of knowledge that would have been acquired had a suitable exemplar been found.

The selection pressure to seek an accessible exemplar is strongest at younger ages, when the knowledge gap between exemplar and learner, *k*(*φ*(*a*))−*k*(*a*), is large so that the cost of a missed learning opportunity is high. As individuals grow older and accumulate knowledge, this gap narrows, and the strength of the selection pressure to seek an accessible exemplar decreases. Consequently, younger individuals are expected to learn preferentially from relatively young exemplars, who are common and still able to provide substantial amounts of novel knowledge, given learners’ limited knowledge at early ages. As individuals age, encountering exemplars with substantially novel knowledge becomes increasingly difficult. Consequently, as they age, individuals are expected to shift toward learning from older individuals, who possess more knowledge. This pattern is observed at ***u***\* (see fig. 2b).

### 3.3 Diversity of evolutionary outcomes: from learning from peers to learning from the oldest

Identifying the factors shaping the evolution of exemplar age choice will help to explain the diversity of age-biased exemplar choice observed in natural populations. Overall, our analyses reveal that these factors generate a wide range of learning strategies, from learners preferentially targeting slightly older age peers to learners consistently learning from the oldest individuals in the population (see fig. 3). We next detail how these factors influence the coevolutionary dynamics of learning-allocation schedules and choice of exemplar age. We summarize the effects of each factor in fig. 3.

**Figure 3:**
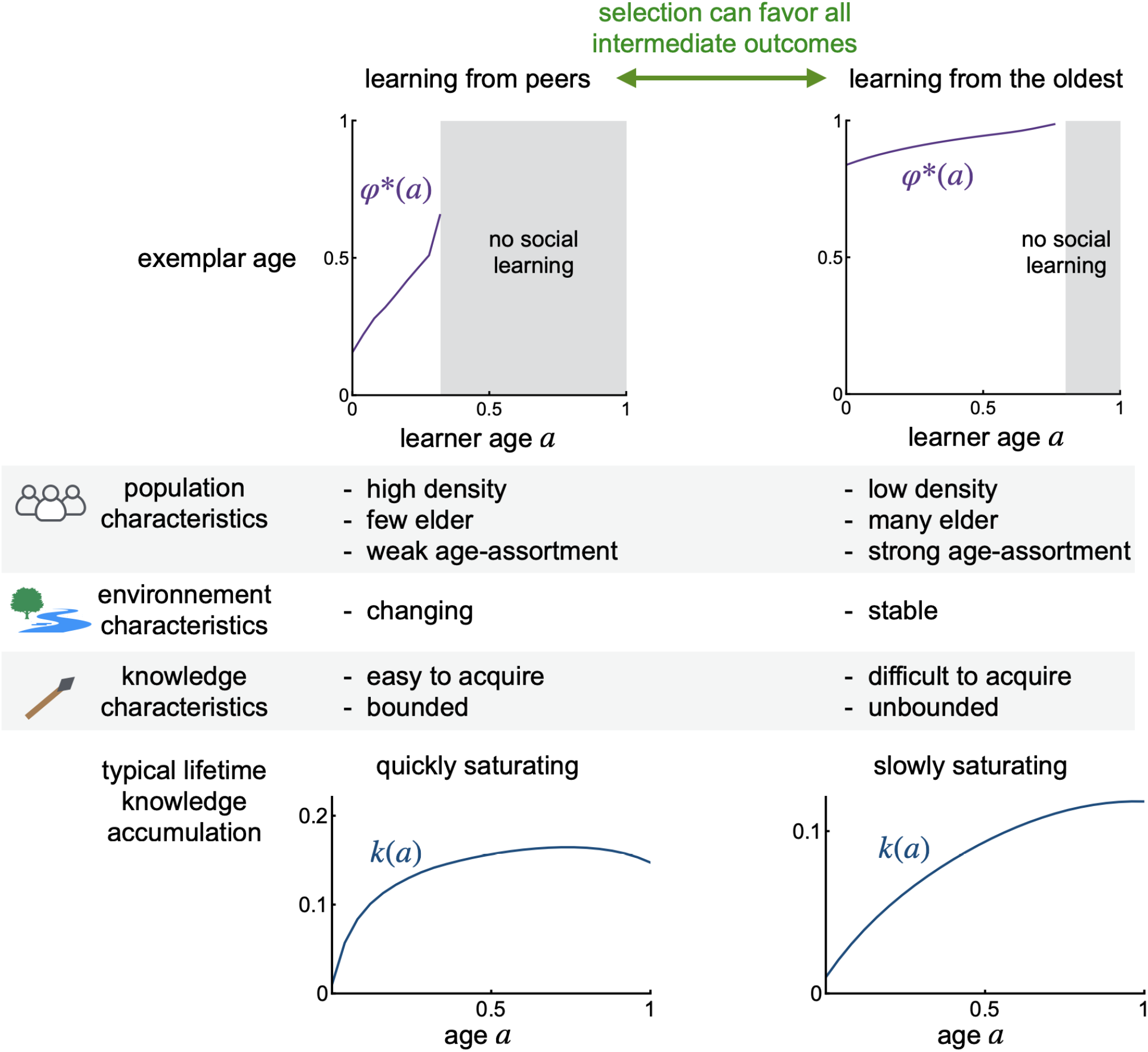
Diversity of evolutionary outcomes in exemplar age choice and their determinants. Top panels show the choice of exemplar age *φ*\*(*a*) at age *a* at ***u***\*, for two contrasting contexts. Depending on population, environment, and knowledge characteristics, selection can favor learning from peers (left) or from the oldest individuals (right), with intermediate outcomes between these extremes also possible. Shaded areas indicate ages at which no social learning occurs. When selection promotes learning from peers, it is generally in cases where lifetime knowledge accumulation saturates rapidly, so that knowledge differences between peers and older individuals remain small. By contrast, when selection promotes learning from the oldest individuals, knowledge accumulation tends to saturate slowly, allowing older individuals to accumulate substantially more knowledge. Parameters for ‘learning from peers’ panel are: *ξ* = 10, *ρ* = 2, *α* = 0.6, *β* = 0.7, *ϵ* = 1, *K* = 1, *µ* = 2. Parameters for ‘learning from the oldest’ panel are: *ξ* = 0.2, *ρ* = 10^−6^, *α* = 0.2, *β* = 0.3, *ϵ* = 0.01, *K* = 10, *µ* = 0.3. Other parameters are: *f* = 100, *σ* = 100, *k*_0_ = 0.01, *e*_min_ = 5, *e*_max_ = 24, *k*_1*/*2_ = 0.2, *M* = 0.01, *γ*_s_ = *γ*_i_ = 0.5, *γ*_b_ = 0.25, *s*_0_ = 0.9, *c* = 10 and *D*(*N*) = 1*/*(1 + 0.1 *N*).

### 3.4 Crowded populations with strong within-cohort interaction structure favor learning from younger exemplars

Here, we examine how population characteristics influence exemplar age choice. Numerical estimates of the identified uninvadable trait schedule vector ***u***\* indicate that, in populations with higher mortality rates *µ*, which consequently contain a lower proportion of older individuals (see figs. S.1a, S.1b), learners at ***u***\* tend to seek younger exemplars (fig. S.1c), as older exemplars are less readily available. In addition, encounters are shaped by population structure, whereby individuals are more likely to encounter others of similar age (e.g., when peers perform similar tasks and interact more frequently). Populations with stronger within-cohort interaction structure (i.e., higher *ρ*) tend to favor the targeting of younger exemplars at ***u***\* (fig. S.2), because learners encounter age peers more frequently.

Other structural factors also shape age preference. In populations with higher encounter rates *ξ* (e.g., denser populations), individuals tend to seek younger exemplars (fig. S.3a). Higher encounter rates increase knowledge transmission, leading to greater knowledge accumulation across the population, such that individuals accumulate more knowledge over their lifetime (fig. S.3b). This amplifies saturation of lifetime knowledge accumulation (fig. S.3b): as individuals accumulate large amounts of knowledge, a greater amount becomes lost through forgetting and obsolescence, making further net knowledge gains increasingly difficult to sustain. Greater saturation then favors learning from younger individuals, as older models offer diminishing access to novel knowledge.

### 3.5 Unstable environments favor learning from younger exemplars

The evolution of exemplar age choice can also be shaped by environmental factors. In environments with higher knowledge loss rates *ϵ* (e.g., more unstable environments with more obsolescence), individuals tend to learn from younger exemplars at ***u***\* (fig. S.4a). Higher loss rates prevent older individuals from accumulating substantial amounts of knowledge (fig. S.4b), reducing the benefit of targeting them as learning exemplars. A high loss rate limits the accumulation of knowledge at the population level, decreasing the amount of knowledge carried by individuals at cultural equilibrium (fig. S.4b). Because less knowledge is available through social transmission, individuals at ***u***\* allocate less time to social learning and more to individual learning (figs. S.4c and S.4d).

### 3.6 Easily acquired knowledge is preferentially learned from younger exemplars

The characteristics of knowledge itself can also influence exemplar age choice. Numerical analyses show that when knowledge is easier to produce (high *α*) and to transmit (high *β*), selection promotes learning from younger individuals at ***u***\* (see figs. S.5a and S.6a). This is because higher rates of knowledge production and transmission promote higher levels of knowledge by increasing its production and transmission (see figs. S.5bc and S.6b). This in turn accelerates the saturation of lifetime knowledge accumulation: as individuals accumulate larger amounts of knowledge, a greater amount becomes lost through forgetting and obsolescence, making further net knowledge gains increasingly difficult to sustain (see figs. S.5bc and S.6b). Consequently, quite young individuals already reach a knowledge level similar to that of older individuals, thereby reducing the advantage of learning from the old.

Interestingly, decreasing the maximum amount of knowledge that can be acquired, *K*, favors learning from younger exemplars at ***u***\* (fig. S.7a). When *K* is small, individuals rapidly approach the maximum attainable knowledge early in life (fig. S.7b). Knowledge therefore saturates early in life, reducing the advantage of targeting older exemplars. However, beyond a certain learner age, most potential exemplars have already reached similar knowledge levels, so only the very oldest individuals retain any remaining knowledge advantage. Low *K* can therefore generate a late shift toward learning from very old exemplars.

## 4 Discussion

We develop a continuous-age model to investigate the evolutionary dynamics of age-specific learning strategies. Our results show that selection favors a progressive reduction in the time allocated to learning with age (fig. 2a). Among the time devoted to learning, individuals rely predominantly on social learning early in life and increasingly on individual learning later on (fig. 2a). This pattern is broadly consistent with previous theoretical work on the evolution of learning schedules (Aoki et al., 2012; Lehmann et al., 2013; Wakano and Miura, 2014). However, these studies either assumed a fixed set of learning stages (Aoki et al., 2012; Wakano and Miura, 2014), or adopted an intermediate framework in which both the durations of learning and resource-exploitation periods and the age-specific allocation between learning modes within a period were allowed to evolve (Lehmann et al., 2013). In contrast to these approaches, our model provides a more biologically realistic framework by allowing individuals to allocate effort among multiple activities at each age (e.g., social learning, individual learning, and energy acquisition), rather than restricting them to a subset of activities at a time. Allowing learning schedules to evolve in this way is important because organisms are unlikely to experience strictly separated learning phases. Instead, learning often continues alongside foraging and reproductive activities throughout life, particularly in long-lived species in which knowledge accumulates gradually over many years (e.g. Whiten and van de Waal, 2018; Musgrave et al., 2021; Pretelli et al., 2022, 2023).

Against this background, our main objective was to investigate how selection shapes the age of social learning exemplars. We show that the evolution of age-biased social learning depends not only on how knowledge changes with age, but also on how accessible individuals of different ages are as exemplars. When knowledge accumulates over the lifespan, older individuals become especially valuable sources of information, and selection can therefore favor learning from them. This interpretation is consistent with evidence that foraging skills, tool use, and other forms of ecological knowledge improve over the lifespan (Musgrave et al., 2021; Pretelli et al., 2022, 2023). However, the most knowledgeable individuals may also be relatively rare or difficult to encounter. Our model shows that this trade-off between exemplar knowledge and accessibility can limit selection for older exemplars and instead favor learning from younger peers, even when those peers are less knowledgeable on average.

We further show that the balance between the benefits of knowledgeable exemplars and the costs of accessing them shifts over the lifespan, leading individuals to preferentially learn from increasingly older exemplars as they age (fig. 2b). This pattern arises because, as learners accumulate knowledge, the pool of individuals possessing knowledge they have not yet acquired becomes progressively smaller. Consequently, the benefits of seeking out the most knowledgeable exemplars increase, favoring a stronger bias toward older individuals. This prediction may help explain recent observations that older chimpanzees exhibit stronger preferences for older social learning exemplars (Nodé-Langlois et al., 2025). More broadly, it suggests that age-dependent social learning biases may emerge as a consequence of learners’ changing knowledge states over the lifespan.

More generally, variation in the balance between exemplar accessibility and knowledgeability may help explain the diversity of age-based social learning strategies observed in nature. Peer learning can evolve despite older individuals being more knowledgeable if their knowledge advantage is outweighed by the costs of accessing them. Our model predicts that differences in population structure, environmental conditions, and patterns of knowledge accumulation can alter the relative benefits of targeting knowledgeable versus accessible exemplars, leading to the evolution of a broad range of age-based learning strategies, from persistent preferences for age peers to strong biases toward the oldest individuals in a population (fig. 3). By linking social learning preferences to demographic structure, environmental conditions, and the distribution of knowledge across age classes, our model provides a framework for understanding when selection should favor different age-based learning biases.

These results also highlight the importance of considering the coevolution of learning schedules and exemplar choice. Deffner and McElreath (2022) showed that unstable environments can favor learning from younger individuals because younger individuals may possess more up-to-date information than older ones. However, this result relies on the assumption that adults cease learning and therefore cannot update their knowledge following environmental change. By contrast, in our model, environmental instability promotes continued investment in learning later in life (fig. S.4cd), allowing older individuals to update their knowledge throughout the lifespan. Consequently, we do not observe conditions under which younger individuals become more knowledgeable than older ones. These results suggest that, when learning schedules are free to evolve, environmental change alone may be insufficient to favor copying peers. Rather, peer learning emerges when the knowledge advantage of older exemplars is outweighed by the costs of locating and interacting with them.

Our formalization nevertheless omits three processes that may refine its predictions. First, age may correlate with knowledge not only because individuals accumulate knowledge, but also because knowledgeable individuals are more likely to survive to older ages (Deffner and McElreath, 2022). Such filtering effects may be especially important when knowledge enhances survival, for example, through predator avoidance or other risk-reducing behaviors (Jeon et al., 2010; Mathiron et al., 2015; Griesser and Suzuki, 2016; Keen et al., 2020; León et al., 2022). Because our model assumes identical knowledge among same-aged individuals and no effect of knowledge on survival, it cannot capture this process. Incorporating knowledge-dependent survival would likely strengthen the age–knowledge association and selection for older exemplars. Second, our model assumes knowledge retains the same value throughout life. In reality, the usefulness of knowledge may vary across life stages, such that younger individuals sometimes possess more relevant knowledge than older individuals. Play provides one example: because play promotes the development of cognitive, motor, and social skills in a relatively low-risk setting, knowledge about appropriate play behaviors may be especially valuable early in life (Bekoff and Byers, 1998; Spinka et al., 2001). Consistent with this idea, human infants preferentially copy peers rather than adults in play contexts (Ryalls et al., 2000). Younger individuals may also be important sources of information when innovations originate disproportionately among them, as documented in Japanese macaques (Kawai, 1965) and capuchins (Perry et al., 2017). Allowing knowledge to peak at different life stages would therefore be a useful extension of the model. Third, we assumed a fixed maximum lifespan beyond which individuals cannot survive. This approximation captures the finite lifespan observed across taxa (Promislow, 1991; Morbey et al., 2005; Bonduriansky and Brassil, 2002) while substantially simplifying the numerical analysis. We do not expect it to affect our qualitative conclusions.

In summary, we show how age-biased social learning can arise from a trade-off between the accessibility and knowledgeability of cultural exemplars. This trade-off emerges from the interplay between demographic structure, learning schedules, and knowledge accumulation, and provides a basis for understanding why individuals learn primarily from peers in some populations, whereas in others they preferentially rely on elders.

## Data and code availability statement

All code used in this study is accessible at github.com/Ludovic-Maisonneuve/age-biased-social-learning.

## Author contributions

LM and LL conceived and designed the study. LM developed the code and performed the analyses. LM wrote the manuscript with contributions from LL.

## Acknowledgments

LM thanks Arthur Weyna for insightful discussions on age-structured models of evolutionary dynamics.

## Appendix A

**Processes underlying knowledge acquisition**

In this section, we derive the rate *A*(*a, φ*_•_(*a*)) at which a focal individual of age *a* with trait schedule vector ***u***_•_ = (*ν*_•_, *s*_•_, *φ*_•_) accepts suitable exemplars (Section A.1) and the rate *r*_il_(*k*_•_(*a*)) of knowledge acquisition during individual learning of such an individual (Section A.2).

### A.1 Derivation of the rate of acceptance of suitable exemplars

To derive *A*(*a, φ*_•_(*a*)) (eq. (3)) recall the assumptions in Section 2.2.1, whereby a focal individual of age *a* encounters other individuals at a rate

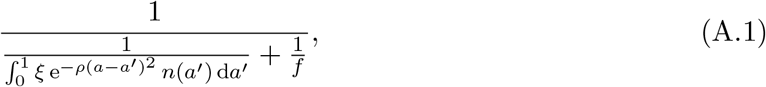

where 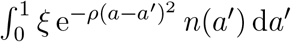 is the total rate at which a focal of age *a* encounters members of the population across all ages *a*^*′*^. Accordingly, 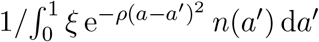 is the mean waiting time until the focal encounters a member of the population, and 1*/f* is the mean duration of an interaction, after which the focal becomes available to encounter another individual.

Given that an encounter has taken place, the probability that the focal individual of age *a*, targeting an exemplar of age *φ*_•_(*a*), accepts the encountered individual as a social learning exemplar is

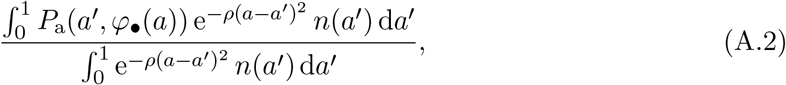

where integrals are taken over all possible ages *a*^*′*^ of encountered individuals.

The rate at which the focal of age *a*, targeting an exemplar of age *φ*_•_(*a*), encounters suitable exemplars is then given by the product of the encounter rate and the probability that an encountered individual is suitable

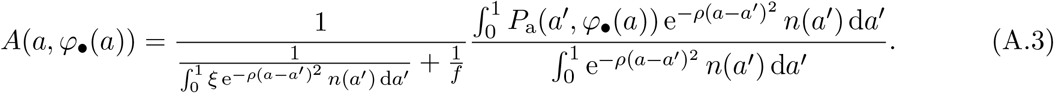

This expression simplifies to

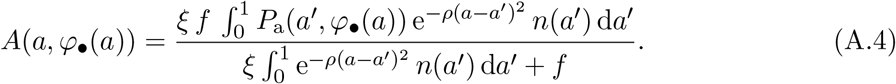

To obtain a simpler expression, we assume that *σ* is sufficiently large that the acceptance kernel 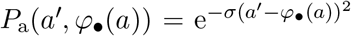 is strongly concentrated around *a*^*′*^ = *φ*_•_(*a*), with characteristic width of order *σ*^−1*/*2^. Over this narrow range, the remaining factor in the integrand, 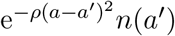 varies slowly with *a*^*′*^. Expanding it around *a*^*′*^ = *φ*_•_(*a*) gives

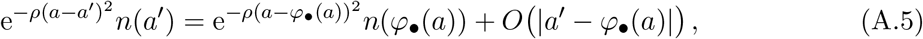

where *O*(*a*^*′*^ − *φ*_•_(*a*)) denotes terms whose magnitude is bounded by |*a*^*′*^ − *φ*_•_(*a*)|.

Since the width of the acceptance kernel is of order *σ*^−1*/*2^, the variation of 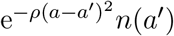 across the region contributing appreciably to the integral is *O*(*σ*^−1*/*2^). Therefore, to leading order,

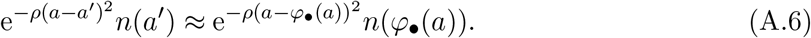

Substituting this approximation into the integral in the numerator of *A*(*a, φ*_•_(*a*)) gives

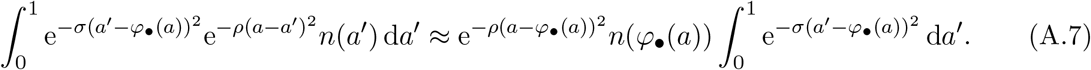

Hence,

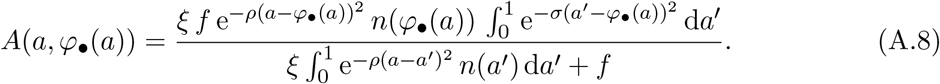

Finally, approximating 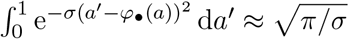 gives

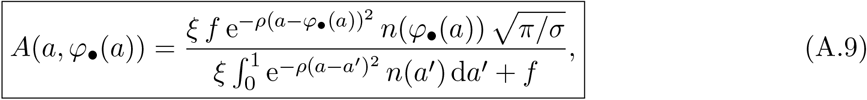

which is eq. (3) of the main text.

### A.2 Rate of knowledge acquisition during individual learning

In this section, we derive eq. (5) of the main text, which gives the rate *r*_il_(*k*_•_(*a*)) at which the focal individual of age *a* with trait schedule vector ***u***_•_ = (*ν*_•_, *s*_•_, *φ*_•_) acquires knowledge under pure individual learning.

We model individual learning as a process of random exploration, in which individuals sample from *I* testable pieces of information at a rate 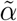, and retain only those belonging to a subset of *K* ⊂ *I* beneficial ones, thereby increasing their knowledge. For example, when attempting to open a nut, *I* includes all possible techniques an individual may try, whereas *K* includes only those that successfully open it. More generally, individuals face a range of challenges (e.g., locating food or recognizing predators), such that *I* represents the set of all testable solutions across these challenges, and *K* the subset that yield successful outcomes. We assume that the ratio *p*_k_ = *K/I* ∈ (0, 1] is constant and treat it like a parameter.

We further assume that individuals do not memorize unsuccessful solutions they have sampled. The probability of discovering a successful solution is given by the ratio of remaining beneficial solutions, *K* − *k*_•_(*a*), to the remaining solutions that can be sampled, *I* − *k*_•_(*a*). Then, under pure individual learning, the focal acquires knowledge at a rate

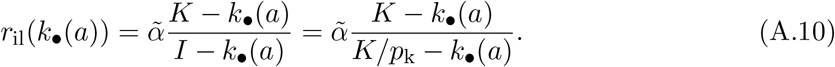

Assuming that most testable pieces of information are not beneficial (i.e., *p*_k_ is small), the rate of knowledge acquisition through individual learning can be approximated by

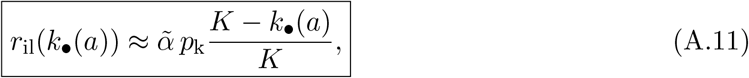

which with 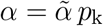 gives eq. (5) of the main text.

## Appendix B

**Resident cultural and demographic equilibrium**

We now characterize the joint cultural and demographic dynamics in a resident population monomorphic at ***u***. For that, we first introduce the coupled dynamics of culture and demographics on the continuous underlying demographic time span. Let then 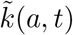 denote the knowledge of a resident individual of age *a* at demographic time *t*, and let *ñ*(*a, t*) denote the density of residents of age *a* at time *t*.

Because individuals age at the same pace as time progresses, the dynamics of 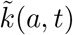 are obtained by substituting 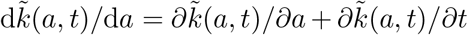 into the left-hand side of eq. (1), and by replacing ***u***_•_ with ***u***, *k*_•_(*a*) with 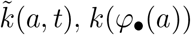 with 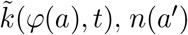 with *ñ*(*a*^*′*^, *t*) and *n*(*φ*_•_(*a*)) with *ñ*(*φ*(*a*), *t*) on the right-hand side of eq. (1) and on eqs. (3) and (4). That is, the subscript _•_ is omitted in all variables appearing in the set of equations (1)–(5) describing the knowledge acquisition process of a focal individual. This yields the partial differential equation

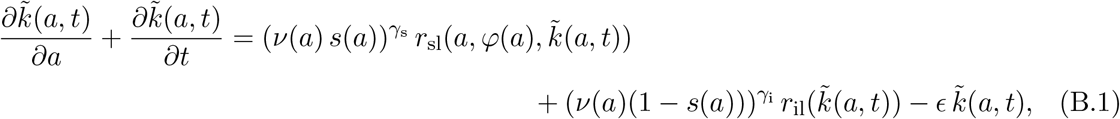

with boundary conditions 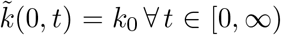 and 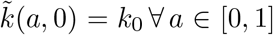, where 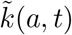 is the solution of eq. (B.1) with the same boundary conditions and where

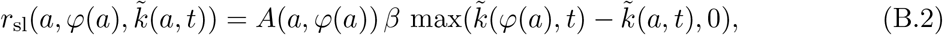

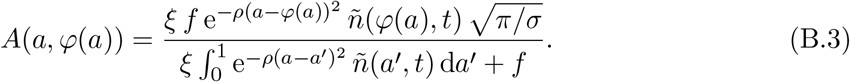

In force of our assumptions of section 2, the demographic dynamics are in turn governed by the McKendrick equation (Keyfitz and Keyfitz, 1997)

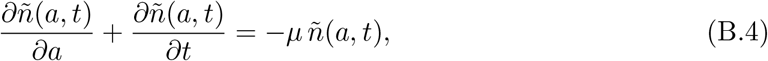

with boundary condition

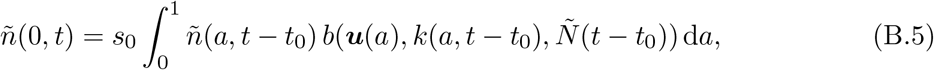

where *t*_0_ is the duration of the juvenile phase and 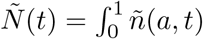 d*a* is the total population size at time *t*. Equation (B.4) describes mortality over the lifespan, while the boundary condition (B.5) captures the density of individuals entering adulthood at time *t*, which equals the number of offspring born at time *t* − *t*_0_ multiplied by the probability *s*_0_ of surviving the juvenile phase to reach maturity.

This coupled dynamical system of equations, eq. (B.1)–(B.5), allows us to evaluate the equilibrium knowledge 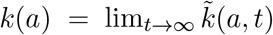 of a resident individual of age *a* in a monomorphic resident population, as well as the density *n*(*a*) = lim_*t*→∞_ *ñ*(*a, t*) of individuals of age *a*, which in turn yields 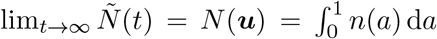. It is not possible to derive analytical expressions for *k*(*a*), *n*(*a*) or *N* (***u***), although note that we have *n*(*a*) = *n*(0)*e*^−*µa*^ and so (B.5) yields at equilibrium the basic reproductive number in the resident population: 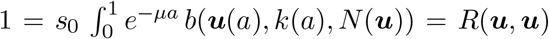. However, all the above quantities can be computed numerically by implementing the joint cultural and demographic dynamics defined by eqs. (B.1)-(B.5) and iterating them until an equilibrium is reached (see Appendix C.3 for details of the procedure). In turn, these expressions allow us to compute the basic reproductive number *R*(***u***_•_, ***u***) (eq. (8)) of a mutant ***u***_•_ introduced as a single copy into the resident population at its cultural-demographic equilibrium.

## Appendix C

**Evolutionary analyses**

In this section, we detail the evolutionary analyses. We first expand the basic reproductive number *R*(***u***_•_, ***u***) of a mutant with trait schedule vector ***u***_•_ to highlight first and second-order selection effects (Section C.1). We then present a representation of the first and second-order age-specific selection pressure (Section C.2). Next, building on the results of the previous sections, we describe the procedure used to numerically approximate a convergence-stable trait schedule vector and to assess its uninvadability (Section C.3). Finally, we analyze the selective pressures acting on learning behaviors and exemplar choice (Section C.4).

### C.1 First and second-order selection

We assume that the mutant trait schedule vector, ***u***_•_, can be written as

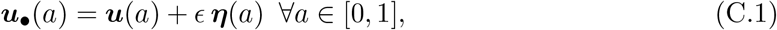

where ***u*** is the resident trait schedule vector, *ϵ >* 0 is small, and ***η*** is a function from [0, 1] to ℝ^3^ such that ***u*** + *ϵ* ***η*** belongs to the space of admissible trait schedule vectors.

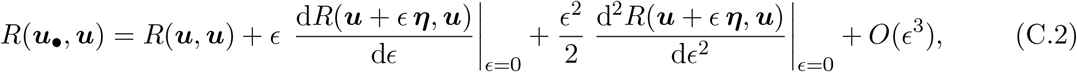

where *R*(***u, u***) = 1 since the resident population is at demographic equilibrium, and *O*(*ϵ*^3^) collects all terms of order *ϵ*^3^ and higher.

#### First-order

Using eq. A.15 in Engel and Dreizler (2013), the first-order derivative of the mutant’s basic reproductive number can be written as an integral over age

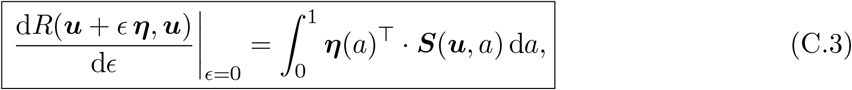

where the age-specific selection gradient ***S***(***u***, *a*) is given by the functional derivative of *R*(***u***_•_, ***u***) at ***u***_•_ = ***u***

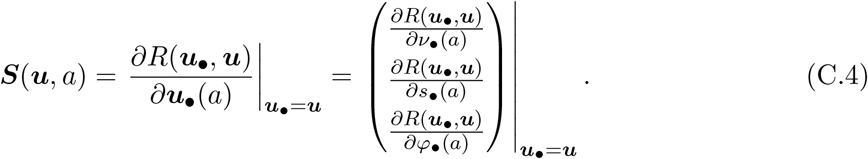

The functional derivative is defined, for a functional *F* of a scalar trait function *u*_1_, by (see eq. A.28 in Engel and Dreizler, 2013)

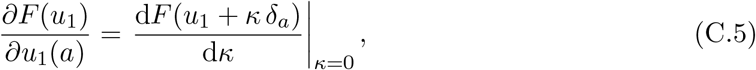

where *δ*_*a*_ is the Dirac delta function centered at *a*, representing a pointwise perturbation of the trait at that age.

#### Second-order

Using eq. A.16 in Engel and Dreizler, 2013, p. 407, the second-order derivative of the mutant’s basic reproductive number can be written as

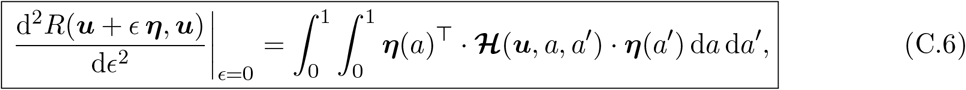

where

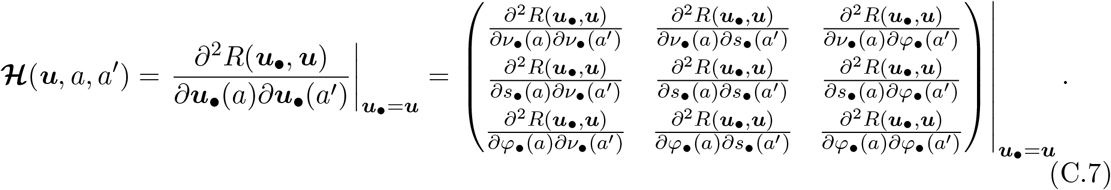

### C.2 Hamiltonian decomposition of first and second-order selection

In this section, we reformulate the first and second-order selection terms to provide a representation of the age-specific selection pressure. To that end, we apply standard computations used in optimal control theory under the calculus of variations approach (e.g., Bryson and Ho, 1975, section 2.3, Kamien and Schwartz, 2012, section 5, Athans and Falb, 2007, section 5.7 for textbook treatments, and Avila et al., 2021, appendix B for previous use in evolutionary biology). These representations then allow us to separate the direct effects of trait changes at a given age from their indirect effects mediated through the internal state dynamics, thereby providing a more tractable expression.

For this purpose, we define the internal state vector of a mutant of age *a* as

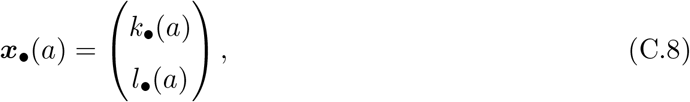

together with the function

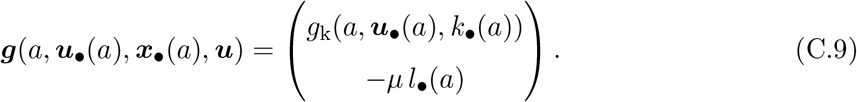

Here, the function *g*_k_(***u***_•_(*a*), *k*_•_(*a*), *k, a*) is given by the right-hand side of eq. (1), and describes the rate of change in mutant knowledge at cultural and demographic equilibrium. The function ***g***(*a*, ***u***_•_(*a*), ***x***_•_(*a*), ***u***) then concatenates the rates of change in knowledge and survivorship of the mutant, so that d***x***_•_(*a*)*/*d*a* = ***g***(***u***_•_(*a*), ***x***_•_(*a*), *u, a*).

As is customary, let us now introduce the Hamiltonian

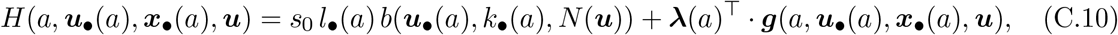

where the costate vector

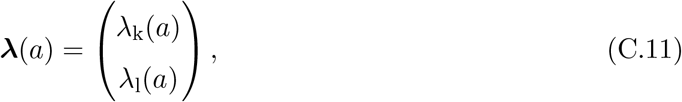

collects the costate variables associated with knowledge and survivorship, respectively; their dynamics are specified below. Here, ^⊤^ denotes the transpose of a vector or a matrix.

We first rewrite the expression of the mutant’s basic reproductive number from eq. (8) by adding the term 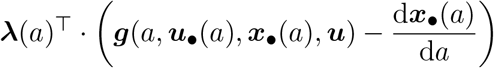 into the integrand. This term is identically zero, since by definition d***x***_•_(*a*)*/*d*a* = ***g***(*a*, ***u***_•_(*a*), ***x***_•_(*a*), ***u***). Thus,

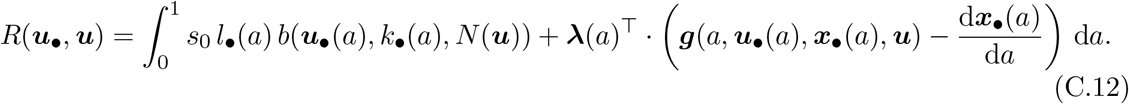

Recognizing the expression of *H*(*a*, ***u***_•_(*a*), ***x***_•_(*a*), ***u***) from eq. (C.10), the mutant basic reproductive number can be expressed as

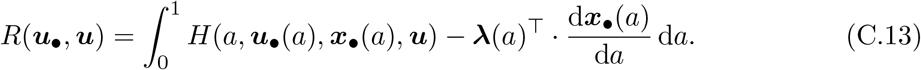

By performing an integration by parts on the second term in the integrand, we obtain

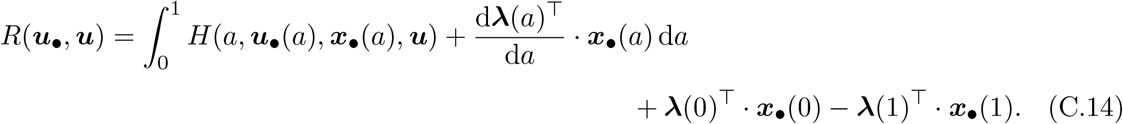

#### First-order

We now compute the age-dependent selection gradient by substituting the expression of *R*(***u***_•_, ***u***) with *a* = *a*^*′*^ from eq. (C.14) into eq. (C.4),

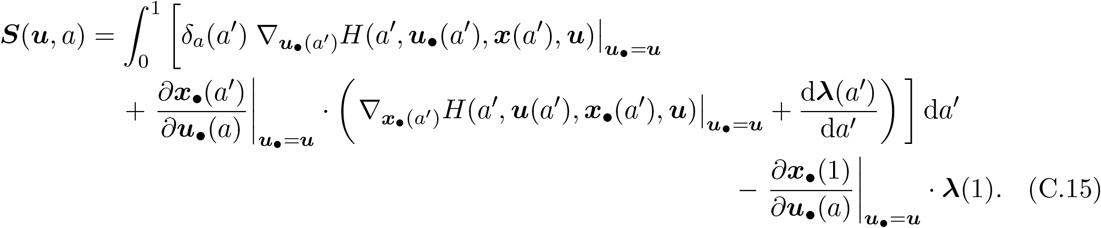

where the operator 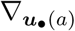 is defined such that for any function *h* of ***u***_•_(*a*)

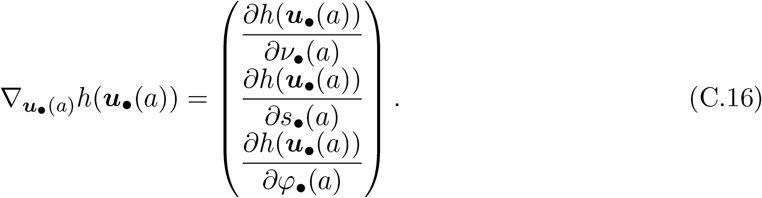

We also adopt the convention that for all *a*^*′*^ ∈ [0, 1]

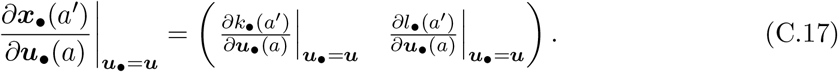

We define the costate vector to satisfy

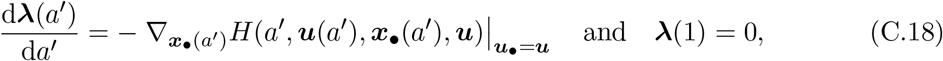

which reduces eq. (C.15) to

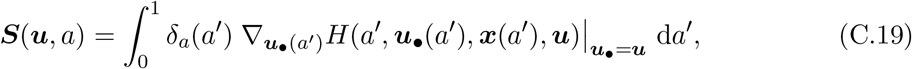

and finally to

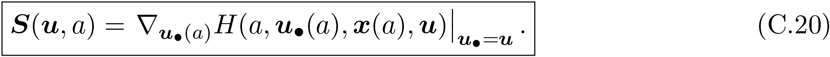

#### Second-order

We now derive the second-order age-dependent selection coefficient. To keep notation concise, we write *H*(*a*) = *H*(*a*, ***u***_•_(*a*), ***x***_•_(*a*), ***u***). Substituting the expression for *R*(***u***_•_, ***u***) given in eq. (C.14) into eq. (C.7) gives

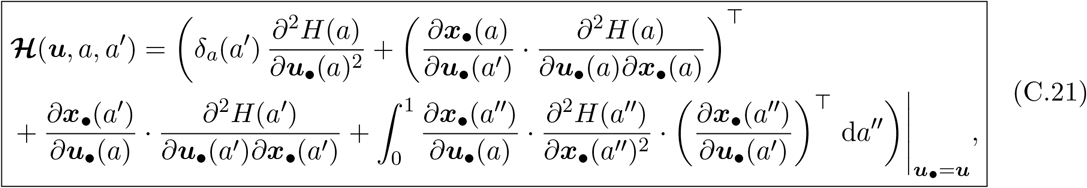

with

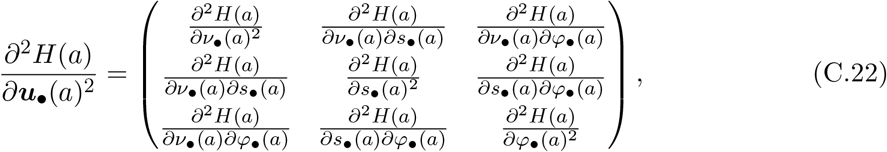

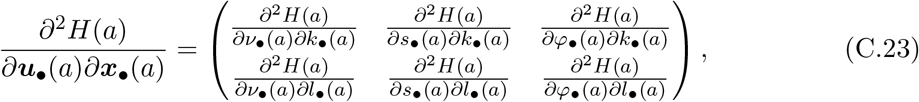

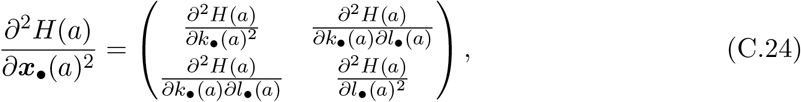

and

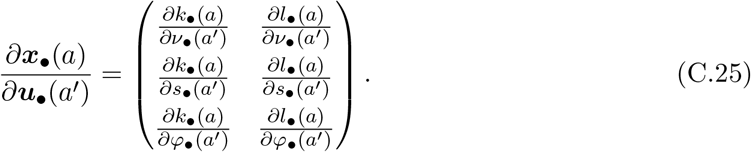

To compute the expression for the second-order selection, we need to derive an expression for 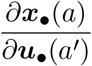 for all *a, a*^*′*^ ∈ [0, 1]. For readability, write ***g***(*a*) = ***g***(*a*, ***u*** (*a*), ***x*** (*a*), ***u***). Using

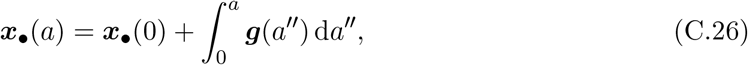

we obtain

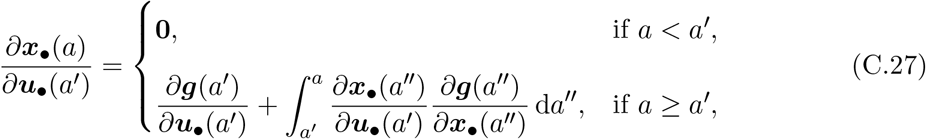

where **0** denotes the zero matrix of appropriate dimension. The resulting ordinary differential equation generally does not admit an analytical solution, but it can be solved numerically.

### C.3 Numerical approximation of an uninvadable trait schedule vector

In this section, we present our approach to numerically approximate a convergence-stable trait schedule vector (Section C.3.1) and to verify its uninvadability (Section C.3.2).

#### C.3.1 Iterative procedure for obtaining a convergence-stable resident trait schedule vector

We describe how to numerically identify a convergence-stable trait schedule vector. We do so by applying the method of steepest descent for functionals (see section ‘The steepest descent algorithm’, pp. 335–336 in Kirk, 2004). We begin by discretizing the age space into small intervals and representing resident trait schedule vectors as piecewise-constant over each subinterval. Let *n*_a_ ≥ 2 denote the number of discrete ages, uniformly distributed over the interval [0, 1], and write 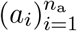 for these discretized ages, with *a*_1_ = 0 and 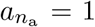. We then introduce an iterative algorithm to identify a convergence-stable trait schedule vector. Let ***u***_*m*_ denote the resident trait schedule vector at iteration *m*. At the initial step (*m* = 1), we set ***u***_1_(*a*) = (0, 0, 1) for all *a* ∈ [0, 1]. The steps of iteration *m* are:

1. *Cultural and demographic equilibrium*. We first numerically compute the equilibrium age-dependent knowledge trajectory in the resident population, *k*(*a*), together with the corresponding equilibrium population size, *N* (***u***_*m*_), at the joint cultural and demographic equilibrium. To this end, we numerically solve the coupled dynamics of the resident knowledge distribution 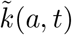 and the age density *ñ*(*a, t*) given in eqs. (B.1) and (B.4) until they reach equilibrium. The coupled dynamical system is solved using a forward finite-difference scheme, with a time step chosen to be half the age step, which enhances numerical stability. The juvenile phase is assumed to last *t*_0_ = 0.1. We assume convergence at time *t*_c_ when

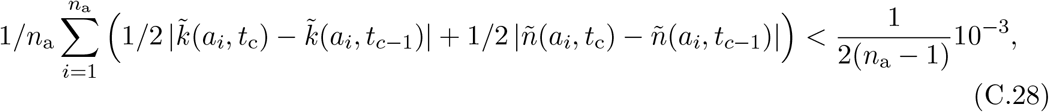

where *t*_*c*−1_ denotes the previous time step in the discretized time grid. We then approximate the resident age-dependent knowledge trajectory by

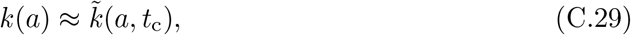

and the equilibrium population size by

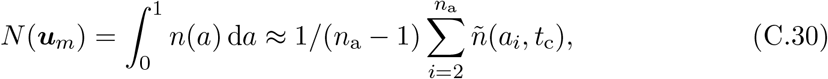

where 1*/*(*n*_a_ − 1) is the width of each age interval.
2. *Costate dynamics*. We compute a numerical approximation of the costate vector function ***λ*** by solving the ordinary differential equation (C.18) with ***u*** = ***u***_*m*_ using the implicit backward Euler method (see Sect. 5.3 p.120 in LeVeque, 2007), using the age discretization mesh 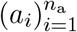. The integration employs the numerical approximations of the function *k* and *N* (***u***_*m*_) obtained in step (i).
3. *Selection gradient*. For each age *a*_*i*_, we compute ***S***(***u***_*m*_, *a*_*i*_) using the expression in eq. (C.20) with ***u*** = ***u***_*m*_. This computation relies on the numerical approximations of the function *k, N* (***u***_*m*_), and the function ***λ*** obtained in steps (i) and (ii).
4. *Update*. We construct the updated resident trait schedule vector ***u***_*m*+1_ as

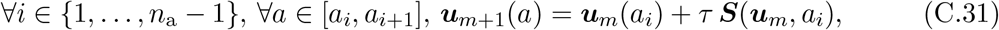

where *τ* is a small step size, and the values of ***S***(***u***_*m*_, *a*_*i*_) are those obtained in step (iii). If necessary, we project ***u***_*m*+1_(*a*) onto the phenotypic space.

The iteration is repeated until 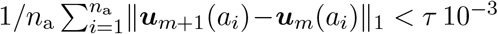, where ∥·∥_1_ denotes the *L*^1^-norm. The vector ***u***_*m*+1_ from the final iteration is then taken as the numerical approximation of an uninvadable resident trait schedule vector, denoted by ***u***\*.

We apply this procedure with *n*_a_ = 26 and *τ* = 0.005.

#### C.3.2 Uninvadability

Once we obtain ***u***\*, we test whether, at a population monomorphic for ***u***\*, selection purges mutations thus promoting monomorphism, or instead allows mutant lineages to invade and coexist, thereby promoting polymorphism. We thus examine whether ***u***\* cannot be invaded by any nearby rare trait schedule vector (small mutation). Here we derive first and second-order conditions for ***u***\* to be uninvadable.

The vector ***u***\* is uninvadable if

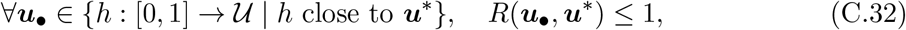

where *{h* : [0, 1] → *U* | *h* close to ***u****}* denotes the set of admissible trait schedule vectors in a neighborhood of ***u***\*, and *U* = [0, 1]^3^ is the set of admissible trait values that an individual may express at each age. Equation (C.32) simply means that, in a population monomorphic for ***u***\*, no sufficiently small admissible mutant trait schedule vector can achieve a basic reproductive number exceeding that of the resident and therefore cannot invade when rare.

##### First-order condition

When the first-order selection is non-null, it dominates the second-order selection, so the success of a mutant lineage is determined by the first-order expansion of *R*(***u***_•_, ***u***\*). Using the expansion of *R*(***u***_•_, ***u***\*) from eq. (C.2) with ***u*** = ***u***\* and *R*(***u***\*, ***u***\*) = 1, retaining terms up to first order, together with the expression of 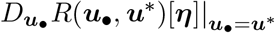 from eq. (C.3) with ***u*** = ***u***\* and ***η***(*a*) = (***u***_•_ − ***u***\*)*/ϵ*, and discretizing, we obtain

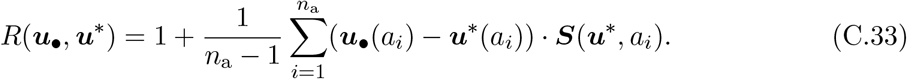

Substituting eq. (C.33) into eq. (C.32) yields the first-order uninvadability condition

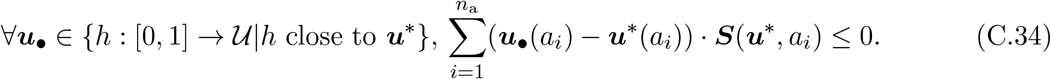

This, in turn, implies that for ***u***\* = (*ν*\*, *s*\*, *φ*\*) to be uninvadable it is necessary that

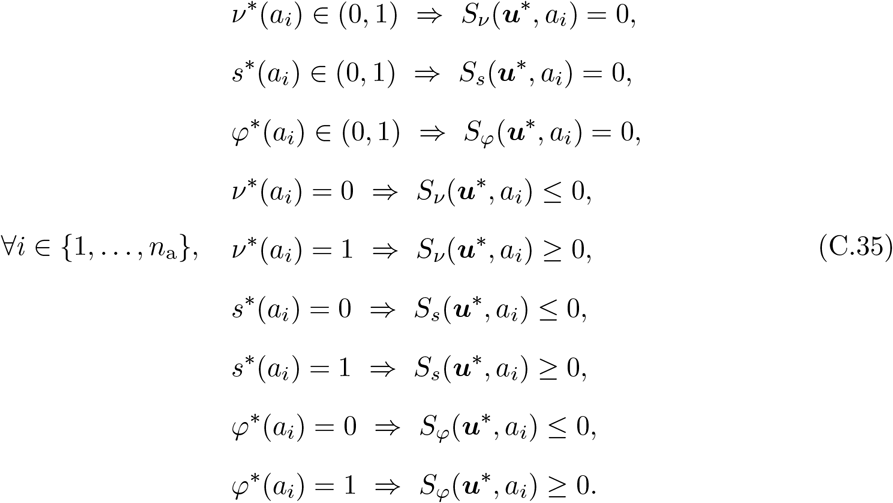

The first three conditions state that if, at a given age *a*_*i*_, a trait lies in the interior of the phenotypic space, then the corresponding component of the selection gradient must vanish at that age. The last six conditions state that if a trait lies on the boundary of the phenotypic space, then no admissible perturbation of that trait at that age can increase mutant fitness. This implies that selection pressures push the trait value expressed at age *a*_*i*_ toward the boundary of the phenotypic space.

Because ***u***\* is obtained by iterating the age-specific selection gradient, it necessarily satisfies the first-order condition (C.35).

##### Second-order condition

The first-order condition (C.35) is necessary but not sufficient for uninvadability: when the first-order term in the expansion of *R*(***u***_•_, ***u***\*) is zero, whether a mutant total reproductive output exceeds that of the resident is determined by the second-order term.

Mutations that shift a trait away from a boundary toward which selection is driving the population cannot invade, as dominant first-order selection acts against them. We therefore restrict attention to first-order neutral mutations, that is, mutations that modify only those trait components at ages for which the corresponding resident component lies in the interior of its feasible set, where first-order selection vanishes. We denote by *ℳ* the set of such mutations. To establish uninvadability, it is thus sufficient to verify that, for all ***u***_•_ ∈ *ℳ* sufficiently small, the second-order variation of *R*(***u***_•_, ***u***\*) is non-positive.

For mutations in *ℳ*, the first-order term vanishes by definition. Dropping the first-order term in eq. (C.2) with ***u*** = ***u***\*, using the expression of d^2^*R*(***u*** + *ϵ* ***η, u***)*/*d*ϵ*^2^ |_*ϵ*=0_ from eq. (C.6) and ***η***(*a*) = (***u***_•_ − ***u***\*)*/ϵ*, and discretizing, we obtain

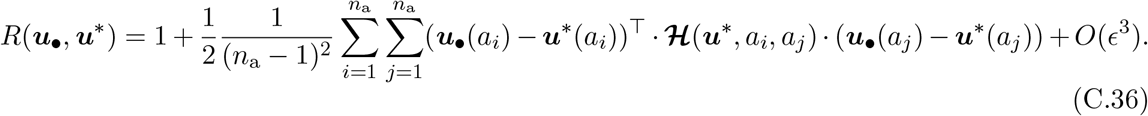

Equivalently, expanding over trait components *i*^*′*^, *j*^*′*^ ∈ *{*1, 2, 3*}*,

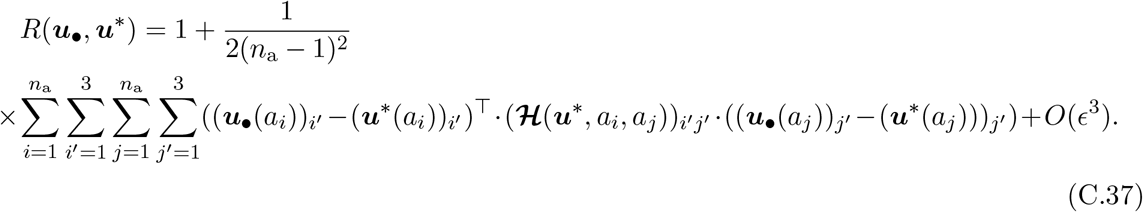

Define the set of interior age–component indices

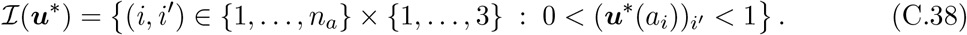

The sums in eq. (C.37) can be restricted to *I*(***u***\*).

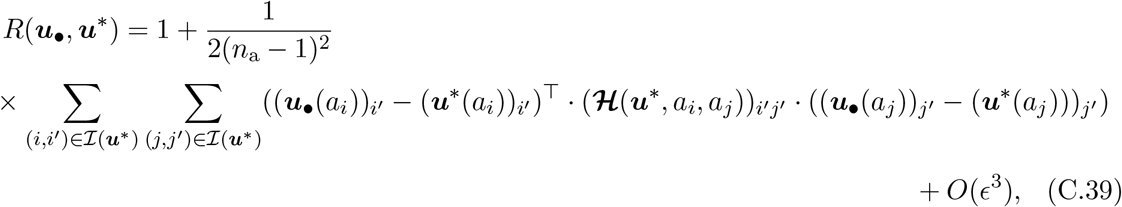

since for any boundary age–component indices (*i, i*^*′*^) ∈*/ ℐ* (***u***\*), we have

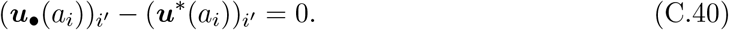

because admissible mutations ***u***_•_ ∈ *ℳ* are allowed to modify only those age–trait components for which the resident value lies strictly between 0 and 1.

Finally, by collecting all interior deviations into a single vector and restricting the Hessian accordingly, we obtain the quadratic form

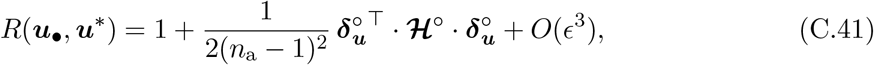

where

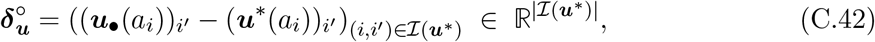

and

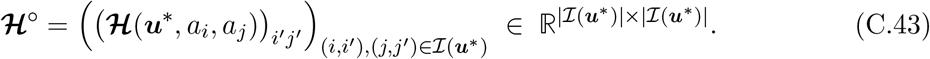

The trait schedule vector ***u***\* is uninvadable if and only if the matrix ***ℋ***^*◦*^ is negative definite, so that the second-order term is strictly negative for every admissible mutation in *ℳ*. In practice, each time we compute ***u***\*, we systematically compute ***ℋ***^*◦*^ and determine whether it is negative definite by checking that all its eigenvalues are strictly negative.

### C.4 Analyzing selection pressures on learning behaviors and exemplar choice

Here, we analyze the selective pressures acting on learning behaviors and exemplar choice. We begin by analyzing the age-specific selection gradient on the allocation of time between learning and energy extraction (Section C.4.1). We then analyze the selection gradient on the allocation of time between social and individual learning (Section C.4.2). Next, we characterize the selection gradient acting on the choice of exemplar age (Section C.4.3), and conclude by detailing how exemplar age choice shapes the probability of encountering a suitable exemplar, thereby clarifying the selective pressures acting on it (Section C.4.4).

#### C.4.1 Analyzing selection pressure on the fraction of time allocated to learning versus reproduction

In this part, we show that the expression of the age-specific selection gradient for *ν*(*a*) reveals that selection typically favors learning in early life and reproduction in later life. To that end, we use the explicit expression for the Hamiltonian *H*(*a*, ***u***_•_(*a*), ***x***_•_(*a*), ***u***), *b*(***u***_•_(*a*), *k*_•_(*a*), *N* (***u***)) obtained from substituting the expression of *b*(***u***_•_(*a*), *k*_•_(*a*), *N* (***u***)) (from eq. (7)), ***λ*** (from eq. (C.11)) and ***g***(*a*, ***u***_•_(*a*), ***x***_•_(*a*), ***u***) (from eq. (C.9) and using the explicit expressions for *g*_k_(***u***_•_(*a*), *k*_•_(*a*), *k, a*), *r*_sl_(*a, φ*_•_(*a*), *k*_•_(*a*)) and *r*_il_(*k*_•_(*a*)) from the right-hand side of eq. (1), eq. (4) and eq. (5)) into eq. (C.10) with ***x***_•_ = ***x***, which gives

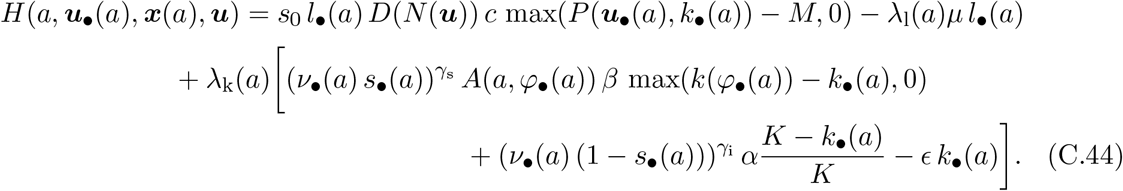

From eq. (C.20) the age-specific selection gradient for *ν*(*a*) is

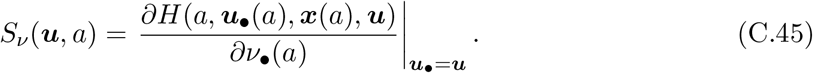

That is, *S*_*ν*_(***u***, *a*) is obtained by taking the partial derivative of the Hamiltonian with respect to *ν*_•_(*a*) while treating the mutant state variables as fixed and equal to the resident values, ***x***_•_(*a*) = ***x***(*a*), and then evaluating the resulting expression at ***u***_•_ = ***u***. Since *ν*_•_(*a*) appears in eq. (C.44) only through the fertility term, via *P* (***u***_•_(*a*), *k*_•_(*a*)), and through the two learning terms, differentiating the Hamiltonian from eq. (C.44) with respect to *ν*_•_(*a*) yields

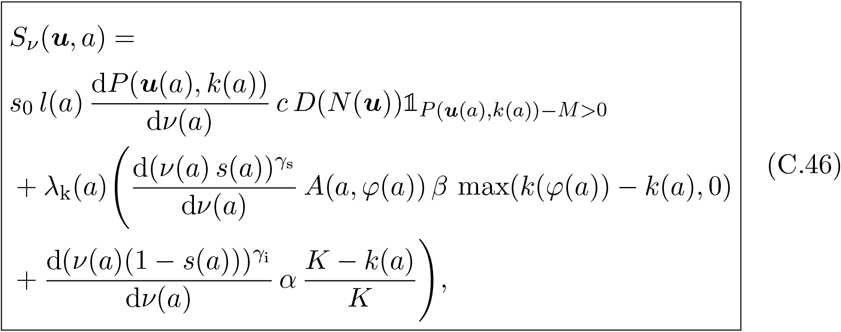

where 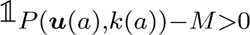 denotes the indicator function, which takes the value 1 when *P* (***u***(*a*), *k*(*a*))− *M >* 0 and 0 otherwise. This term arises from differentiating the function max(*P* (***u***(*a*), *k*(*a*)) − *M*, 0) with respect to *ν*(*a*).

The first term in eq. (C.46) is the fecundity cost of allocating time to learning rather than to reproduction. The two terms on the second line capture the benefits to the remaining reproductive output from increased knowledge gained through social and individual learning, respectively. These benefits are proportional to the costate variable *λ*_k_(*a*), which measures how a change in knowledge at age *a* affects the remaining reproductive output. The costate variable *λ*_k_ typically declines with age, as the amount of remaining reproduction decreases (fig. C.1). Consequently, the benefit of learning should typically decrease with age. In addition to this life-history effect, the benefits of both social and individual learning further decrease as individuals accumulate knowledge. Altogether, these effects suggest that selection favors an allocation of time to learning *ν*(*a*) that peaks after birth (i.e., *a* = 0) and declines with age.

**Figure C1:**
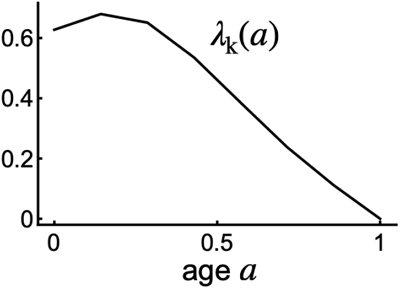
Costate variable *λ*_k_(*a*) as a function of age *a* corresponding to fig. 2 in the main text.

#### C.4.2 Analyzing selection pressures on the fraction of learning time allocated to social versus individual learning

In this part, we show that the expression of the age-specific selection gradient for *s*(*a*) reveals that, among the time allocated to learning, selection typically favors social learning early in life and individual learning later in life.

Differentiating the Hamiltonian in eq. (C.44) with respect to *s*_•_(*a*), and then evaluating the resulting expression at ***u***_•_ = ***u***, following the same steps as above, yields

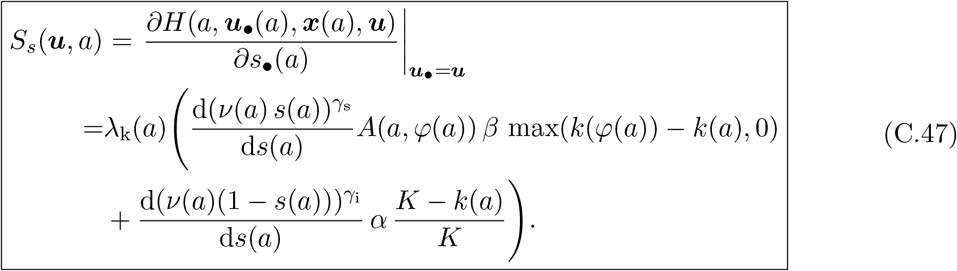

The two terms in brackets in eq. (C.47) are, respectively, the benefit to remaining reproductive output of increasing social learning and the cost of reducing individual learning. The benefit of social learning declines sharply with age because the knowledge gap between a learner and their exemplar, *k*(*φ*(*a*)) − *k*(*a*), narrows rapidly over time. In contrast, the potential gains from individual learning decrease much more slowly, since *K > k*(*φ*(*a*)). As a result, the relative benefit of allocating time to individual rather than social learning generally increases with age, leading selection to favor a shift toward individual learning over the life course (i.e., a decrease in *s*(*a*)).

#### C.4.3 Selection pressures on the age choice of the learning exemplar

Here we derive eq. (10) of the main manuscript, which provides the term to which the age-specific selection gradient for *φ*(*a*) is proportional.

Differentiating the Hamiltonian in eq. (C.44) with respect to *φ*_•_(*a*), and then evaluating the resulting expression at ***u***_•_ = ***u***, following the same steps as above, yields

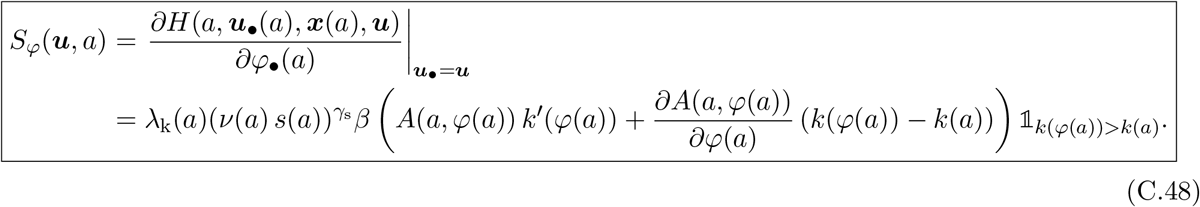

Equation (C.48) corresponds to eq. (10) in the main text, up to a proportionality factor that does not affect the sign of selection.

#### C.4.4 Impact of age choice of exemplar on the probability of finding a suitable exemplar

Here we derive eq. (11) of the main manuscript, which provides the term to which *∂A*(*a, φ*(*a*))*/∂φ*(*a*) is proportional.

Using the expression of *A*(*a, φ*(*a*)) from eq. (3), yields

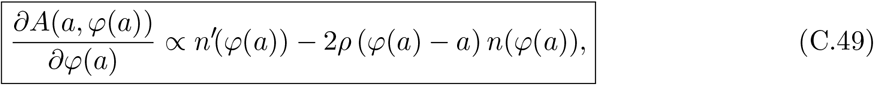

which is eq. (11) in the main text.

Then we show that *n*^*′*^(*φ*(*a*)) is negative. Taking eq. (B.4), evaluated at *a* = *φ*(*a*), at demographic equilibrium, where lim_*t*→∞_ *n*(*φ*(*a*), *t*) = *n*(*φ*(*a*)), so that *∂n*(*φ*(*a*), *t*)*/∂a* = *n*^*′*^(*φ*(*a*)), *∂n*(*φ*(*a*), *t*)*/∂t* = 0, and *n*(*φ*(*a*), *t*) = *n*(*φ*(*a*)), yields

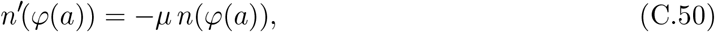

which implies that *n*^*′*^(*φ*(*a*)) *<* 0.

## D Supplementary figures

**Figure S.1:**
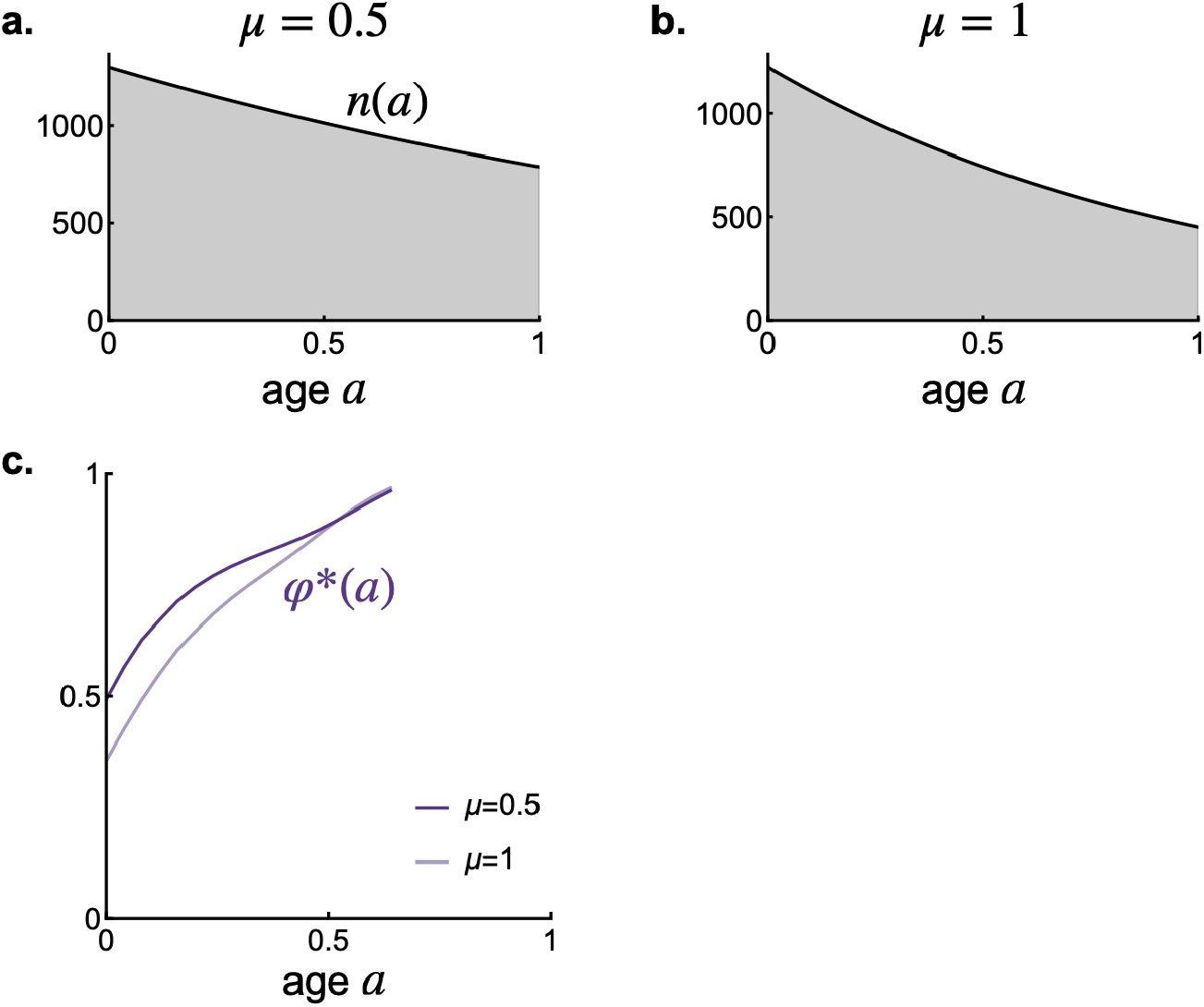
Effect of *µ* on age-specific density and choice of exemplar age. Age-specific density for **a** *µ* = 0.1 and **b** *µ* = 1. **c** Choice of exemplar age, *φ*\*(*a*), by a learner of age *a* at ***u***\* for *µ* = 0.1 and *µ* = 1. Default parameter values (used throughout all supplementary figures unless stated otherwise): *ξ* = 1, *f* = 100, *ρ* = 0.01, *σ* = 100, *k*_0_ = 0.01, *α* = 0.5, *β* = 0.6, *ϵ* = 0.1, *K* = 10, *µ* = 1, *e*_min_ = 5, *e*_max_ = 24, *k*_1*/*2_ = 0.2, *M* = 0.01, *γ*_s_ = *γ*_i_ = 0.5, *γ*_b_ = 0.25, *s*_0_ = 0.9, *c* = 10 and *D*(*N*) = 1*/*(1 + 0.1 *N*).

**Figure S.2:**
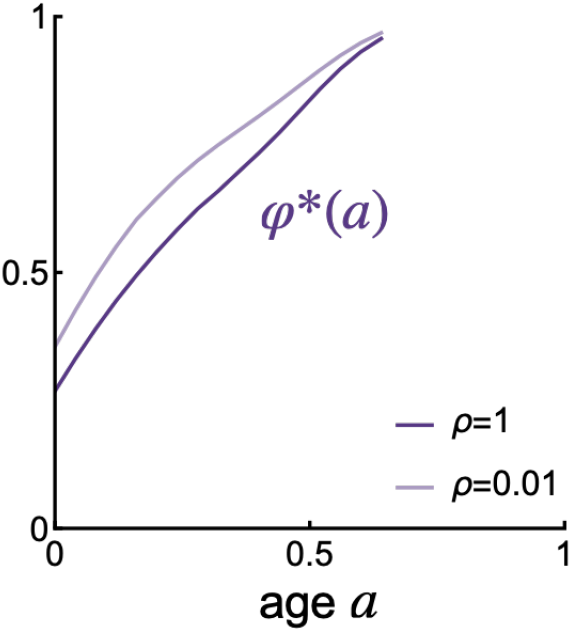
Effect of *ρ* on choice of exemplar age. Choice of exemplar age, *φ*\*(*a*), by a learner of age *a* at ***u***\* for *ρ* = 0.01 and *ρ* = 1. Default parameters are given in fig. S.1.

**Figure S.3:**
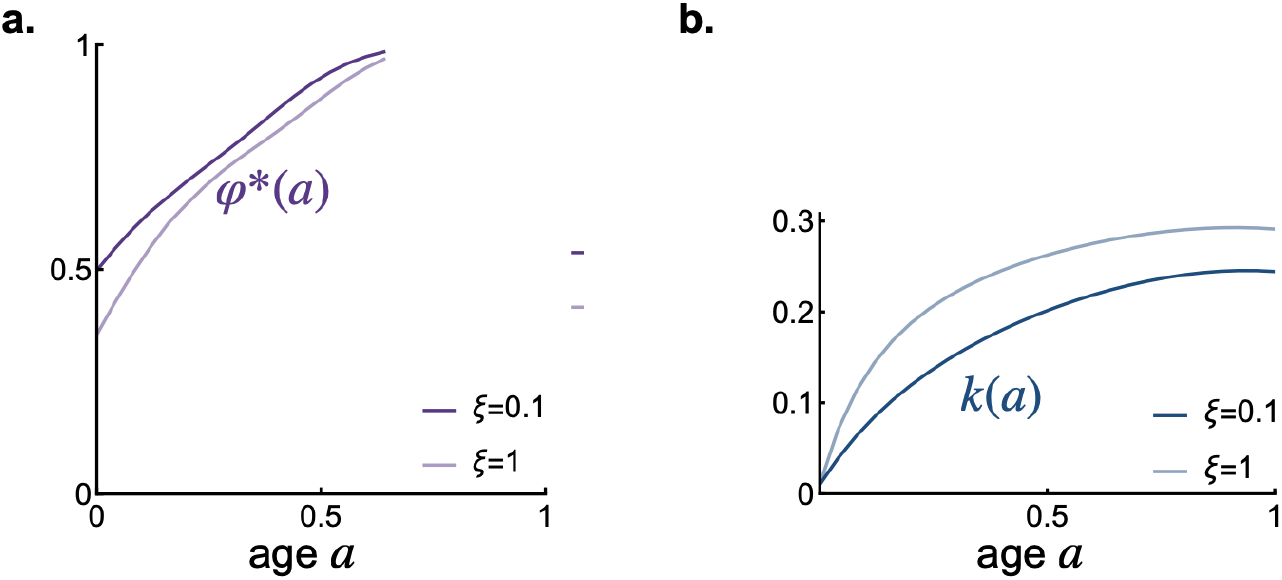
Effect of *ξ* on choice of exemplar age and lifetime knowledge accumulation. **a** Choice of exemplar age, *φ*\*(*a*), **b** knowledge acquired, by a learner of age *a* at ***u***\* for *ξ* = 0.1 and *ξ* = 1. Default parameters are given in fig. S.1.

**Figure S.4:**
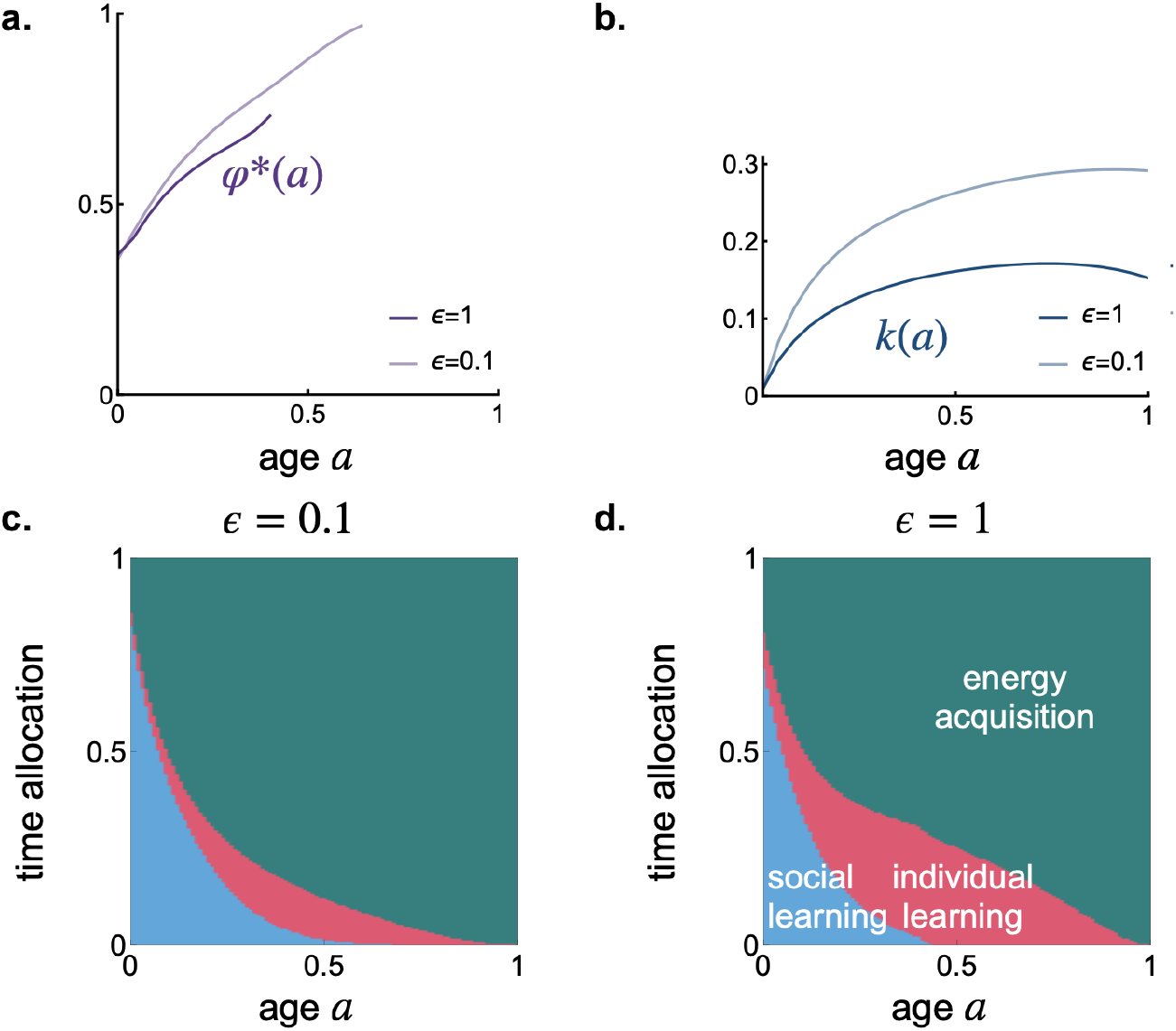
Effect of *ϵ* on choice of exemplar age, lifetime knowledge accumulation, and time allocation schedule. **a** Choice of exemplar age, *φ*\*(*a*), **b** knowledge acquired, by a learner of age *a* at ***u***\* for *ϵ* = 0.1 and *ϵ* = 1. Time allocation schedule at ***u***\* for **c** *ϵ* = 0.1 and **d** *ϵ* = 1 (plot details are provided in the legend of fig. 2). Default parameters are the same as in fig. S.1.

**Figure S.5:**
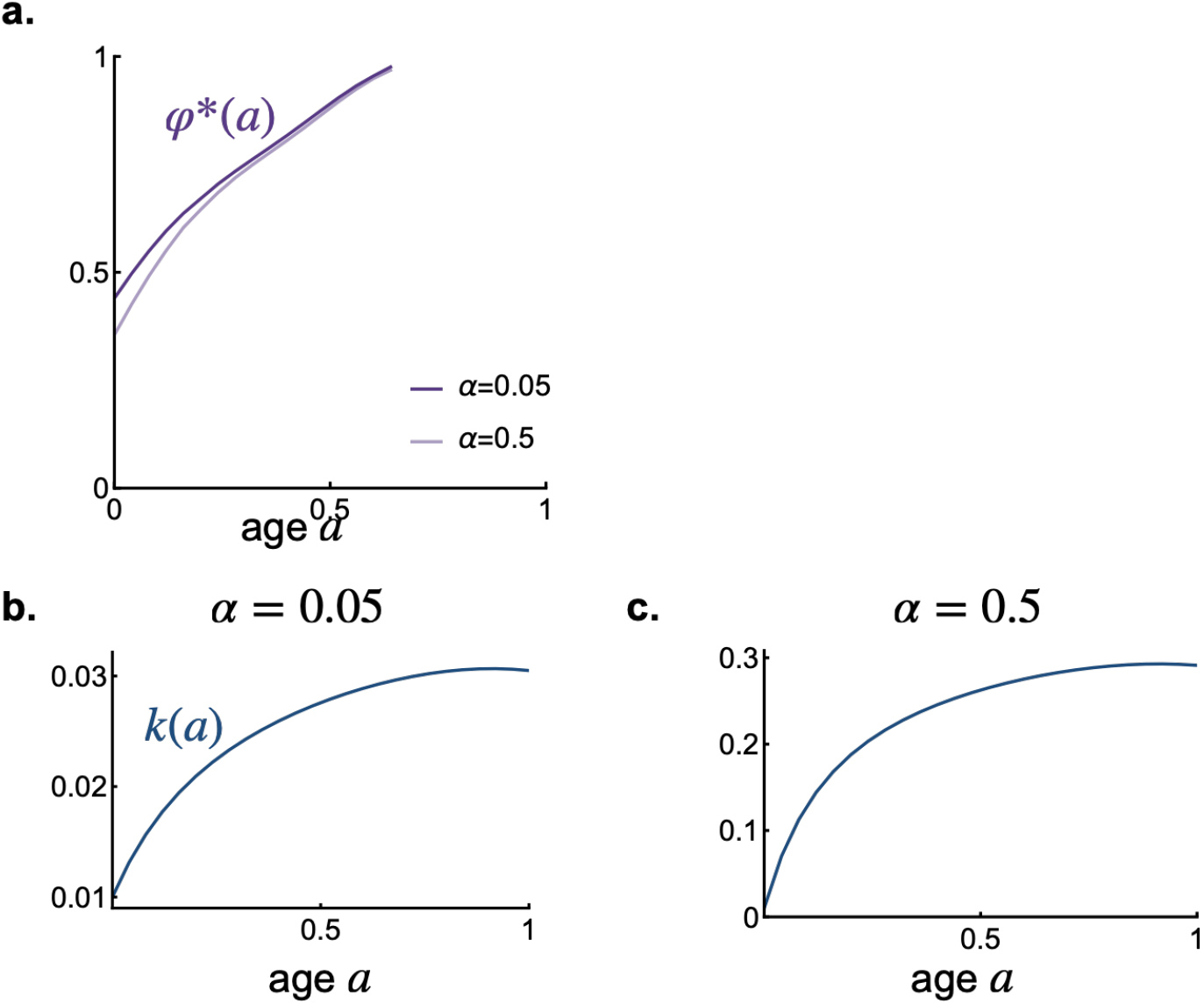
Effect of *α* on choice of exemplar age and lifetime knowledge accumulation. **a** Choice of exemplar age, *φ*\*(*a*) at ***u***\* for *α* = 0.05 and *α* = 0.5. Knowledge acquired, by a learner of age *a* at ***u***\* for **b** *α* = 0.05 and **c** *α* = 0.5. Default parameters are given in fig. S.1.

**Figure S.6:**
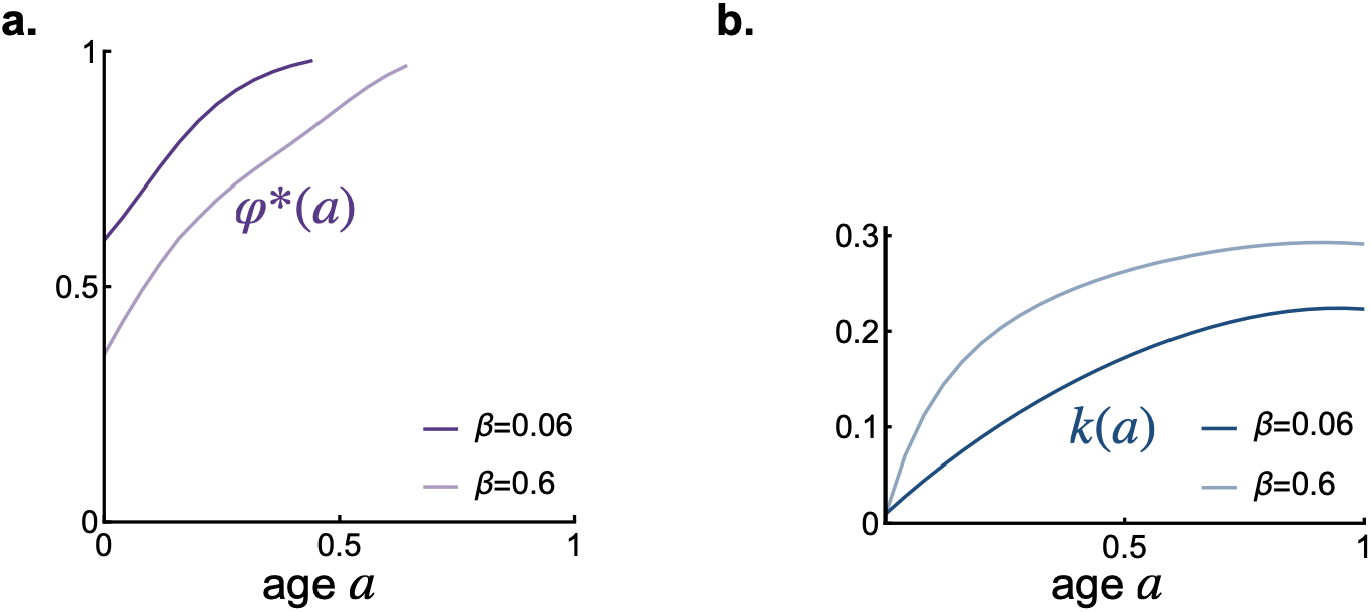
Effect of *β* on choice of exemplar age and lifetime knowledge accumulation. **a** Choice of exemplar age, *φ*\*(*a*), **b** knowledge acquired, by a learner of age *a* at ***u***\* for *β* = 0.06 and *β* = 0.6. Default parameters are given in fig. S.1.

**Figure S.7:**
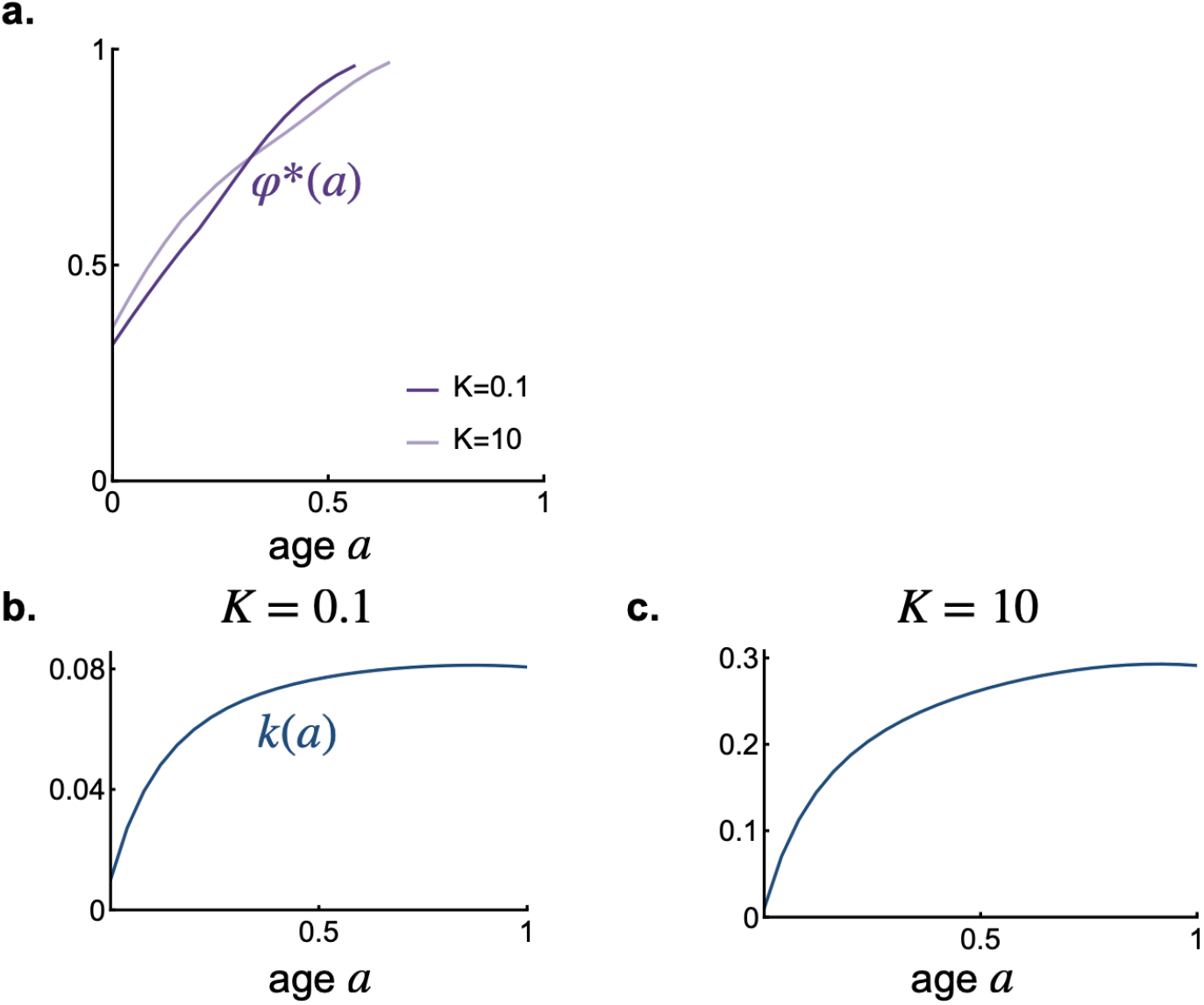
Effect of *K* on choice of exemplar age and lifetime knowledge accumulation. **a** Choice of exemplar age, *φ*\*(*a*) at ***u***\* for *K* = 0.1 and *K* = 10. Knowledge acquired, by a learner of age *a* at ***u***\* for **b** *K* = 0.1 and **c** *K* = 10. Default parameters are given in fig. S.1.

